# *Stenocytosis*: a mechanism for supramolecular attack particle transfer at the CTL lytic synapse

**DOI:** 10.1101/2025.11.05.686769

**Authors:** Lucie Demeersseman, Claudia Schirra, Ute Becherer, Marie-Pierre Puissegur-Lay, Brienne McKenzie, Michael L. Dustin, Sabina Müller, Salvatore Valitutti

## Abstract

Supramolecular attack particles (SMAPs) are recently discovered key components of the cytotoxic T lymphocytes (CTLs) lytic arsenal exhibiting autonomous cytotoxic behavior. Yet, how these particles might serve as synaptic weapons transferred from CTL into target cells within dynamic lytic synapses remains to be elucidated.

In CTL interacting with immobilized stimuli, total internal reflection fluorescence microscopy (TIRFM) and rapid live 3D cell imaging showed that granules containing SMAPs navigate through narrow cortical actin cytoskeleton depletion areas to reach plasma membrane secretion hot-spots where SMAPs are released. In CTLs interacting with cognate target cells, correlative light-electron microscopy (CLEM) and structured illumination 3D imaging showed SMAP release into the synaptic cleft and penetration into the target cell trough an equally narrow gap in the target cell cortical actin cytoskeleton mirroring that formed in the CTL cytoskeleton, and just large enough to admit the SMAP into an early endosome.

Our results reveal a previously undescribed process, *stenocytosis* (from the ancient Greek στενός, “narrow”), which allows hundred-nanometer-scale lytic particles to exit CTLs and enter target cells while evading synaptic defenses.

## Introduction

Many immunotherapeutic strategies against cancer (including immune checkpoint blockade, bispecific antibodies, CAR-T cells, and others) rely upon successful deployment of cytotoxic T lymphocytes (CTLs) to eliminate tumor cells in an antigen-specific manner. CTLs recognize their targets through engagement of their T-cell receptors (TCRs) by peptide MHC (pMHC) class I complexes displayed on the target cell surface. Upon productive TCR engagement, CTLs increase their adhesion with target cells, arm their lytic machinery and re-direct the cytotoxic arsenal towards target cells to provide lethal hits within specialized contact areas named *lytic synapses*. To kill target cells, CTLs are endowed with an array of lytic molecules including perforin, granzymes, and death receptor ligands ((*1*) (*2*)). Additionally, they produce cytokines known to influence cytotoxic behavior, such as IFNγ and TNFα ((*1*) (*2*)). CTLs can also release supramolecular attack particles (SMAPs), autonomous killing entities containing lytic molecules enrobed by a glycoprotein shell (*3*) (*4*). These particles can act as “cytotoxic bombs” to kill target cells from a distance (*3*) (*5*). Ultimately, these various killing strategies converge upon the engagement of the target cell’s endogenous executioner proteins (e.g. caspases) and the commitment of the target cell to programmed cell death (PCD).

A peculiar characteristic of CTL responses is their exquisite sensitivity to antigenic stimulation, and indeed CTLs can be triggered to lytic granule secretion by a very small number of pMHC complexes on the target cell surface ((*1, 6, 7*)). CTLs are also extremely rapid in annihilating their targets, secreting soluble perforin and granzymes within seconds after TCR triggering ((*1, 8*)). Finally, CTLs can engage several target cells simultaneously or sequentially and induce cytotoxicity through multiple rounds of lethal hit delivery ((*9*) (*10, 11*)). Despite the remarkable rapidity and sensitivity of CTL attack, and the breadth of cytotoxic molecules available for deployment, we and others, have shown that CTL-mediated cytotoxicity is limited by the ability of cancer cells to engage ultra-rapid defense mechanisms (e.g. membrane repair) at the lytic synapse ((*12*) (*13*) (*14*)). These findings positioned the lytic synapse as a key battleground within which pro-death and pro-survival signals compete to determine target cell fate.

The possibility that SMAPs might serve as autonomous killing entities has raised interest in these lytic particle as potential cell-free weapons able to fight cancer by circumventing tumor cell resistance at the lytic synapse ((*3*)). Nevertheless, whether SMAPs might also serve as synaptic weapons and contribute to some steps of target cell annihilation at the CTL/target cell contact site remains to be elucidated.

In this study we investigated the contribution of SMAPs in the paradigm of lethal hit delivery to target cells at the lytic synapse. In particular, we addressed the unresolved question of whether and how hundred nanometer sized lytic particles might exit CTLs and penetrate into target cells within the context of dynamic cell-cell contacts.

To address this and related questions, we employed a combination of high-resolution photonic and electronic microscopy imaging techniques to illustrate the routing of SMAPs from CTLs lytic machinery into cognate target cells via the synaptic cleft.

Our study reveals that CTLs deployed SMAPs as rapid and directionally secreted synaptic weapons during the early steps of CTL/target cell interaction.

In CTLs triggered by immobilized stimuli, TIRFM and rapid live 3D cell imaging showed that SMAPs percolated through areas of CTL cortical actin cytoskeleton depletion and underwent a swift secretion process via the formation of short-lived docking and fusion events in the plasma membrane. In CTLs conjugated with cognate target cells, structured illumination 3D imaging and CLEM showed that SMAPs were secreted into the synaptic cleft within minutes after cell-cell contact. Secreted particles penetrated target cells through narrow cortical actin cytoskeleton depletion areas that mirrored those formed in CTL cytoskeleton, revealing a previously unidentified mechanism of cell-cell supramolecular particle transfer called *stenocytosis* (from the ancient Greek στενός, narrow). SMAPs were found in the target cell endocytic compartment, suggesting that the previously described process of target cell membrane reparation triggered by perforin might favor SMAP uptake ((*13*) (*15*)). All in all, our results indicate that SMAPs might constitute a second line of synaptic weapons that penetrates into target cells despite membrane reparation.

## Results

### SMAP release hot-spots are detected within minutes after TCR engagement

To study the potential role of SMAPs as synaptic weapons, we initially assessed the spatio-temporal characteristics of their secretion following TCR engagement. To this end, we took advantage of the notion that SMAPs can be intracellularly traced by an overnight incubation of human CD8^+^ T cells (polyclonal T cell lines or antigen specific T cell clones) with fluorochrome-labelled wheat germ agglutinin (WGA) that binds the SMAP glycoprotein shell ((*3*) (*4*)). Correlative light and electron microscopy (CLEM) showed that WGA staining overlaps with multi core granules (MCG) a distinct class of cytotoxic granules containing SMAPs ((*4*)). Moreover, in our clonal CTLs, WGA^+^ SMAP dimension correlated with dimensions previously measured in human and mouse CTL ((*3*) (*4*) (*16*)). Together, these observations validated the use of this dye for specific SMAP labelling in our cellular model (**Supplementary Fig. 1**).

To study the dynamics of SMAP release we imaged human CTL clones or polyclonal CD8^+^ T cells (*in vitro* expanded from peripheral blood) using TIRFM. Cells were pre-loaded with WGA-AF647 and stained with Vybrant™ Dil to visualize cell membrane, seeded on µ-slide chambers (ibidi) previously coated with anti-CD3/ICAM-1 or ICAM-1 only and imaged at 37°C. This approach showed that TCR/CD3-LFA-1 engagement triggered the formation of distinct areas of SMAP release hot-spots, characterized by the accumulation of hundred nanometers sized WGA^+^ particles within domains exhibiting significantly reduced plasma membrane staining (**Fig. 1a**, **Supplementary Movie 1-3**). This process was detected within minutes after T cell contact with the stimulating surface and was absent in cells stimulated with immobilized ICAM-1 only exhibiting uniform Vybrant™ Dil staining (**Supplementary Movie 4**).

**Figure 1.**
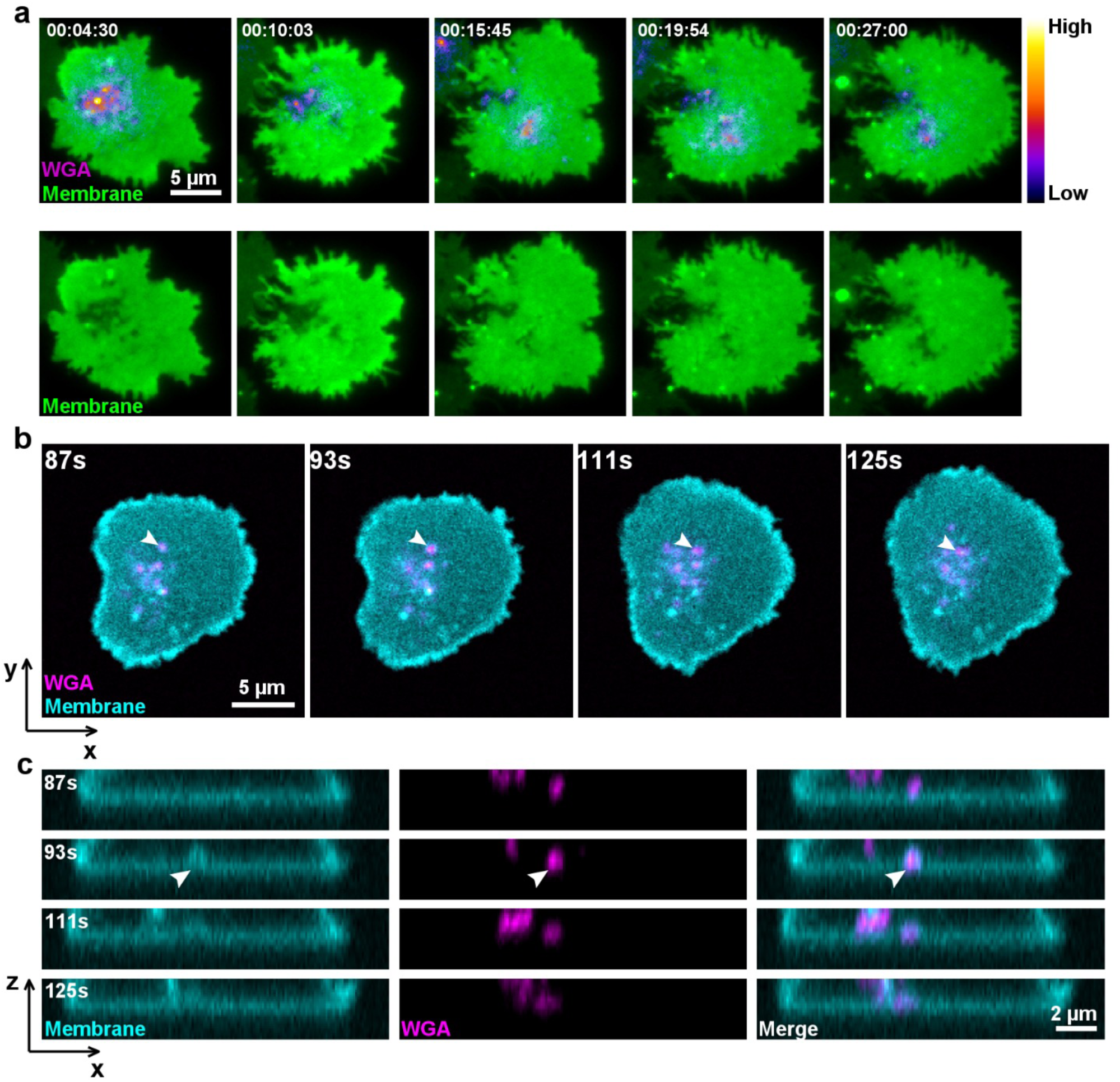
SMAP release hot-spots are detected within minutes after TCR engagement. **(a)** Live TIRF imaging of CTL seeded on CD3/ICAM-1 coated ibidi µ-slide chambers 37°C/5%CO_2_. CTLs were pre-loaded with WGA-AF647 (pseudocolor) and stained with Vibrant Dil (green). Still images are from **Supplementary Movie 1-3**. Upper panel: appearance of WGA^+^ particles in the TIRF plane (**Supplementary Movie 1)** Lower panel: Corresponding domains exhibiting reduced plasma membrane staining (**Supplementary Movie 3**). Data are from one representative experiment out 4. **(b)** Ultra-rapid spinning-disk 3D imaging of a representative CTL pre-loaded with WGA-AF647 (magenta) and Vibrant^TM^ DiD (cyan) seeded on CD3/ICAM-1 coated ibidi µ-slide chambers. Panels show x/y still images from live cell acquisition. **(c)** x/z axis images of the same cell. Still images are from live z-stack acquisition. Left panels: vesicle fusion (cyan); middle panels: WGA^+^ particles (magenta) reaching the 3D area of inspection; right panels: WGA^+^ particle secretion in concomitance with vesicular fusion events. Images are from **Supplementary Movie 5-6**. Data are from one representative experiment out 3.

To define whether the reduced plasma membrane staining upon TCR stimulation was due the intense endo/exocytic process which is known to take place in CTLs during lytic granule secretion and TCR internalization ((*13*) (*17*)), we closely inspected the surrogate synaptic area using 3D high spatio/temporal resolution imaging. Cells, pre-loaded with WGA-AF488, were stained with Vybrant™ DiD and the narrow area of contact between CTL and stimulating surface was inspected by rapid Z-stacks acquisition using a spinning-disk microscope at 37°C. As show, in **Figure 1b-c** and **Supplementary Movie 5 and 6**, this experimental approach highlighted rapid docking and fusion events of vesicles carrying WGA^+^ particles, reminiscent of Ω-profile merging described in neuro-endocrine cells ((*18*) (*19*) (*20*)).

Taken together, these observations indicate that human CTLs contain stocks of pre-formed SMAPs that are rapidly released after TCR triggering within secretion hot-spots characterized by enhanced membrane turnover.

### Secreted SMAPs percolate through the CTL actin cytoskeleton cortical network

The above results raised the question if the observed hot-spots of SMAP release were accompanied by re-modeling of the sub-membrane actin cytoskeleton network, to allow the passage of MCGs containing SMAPs ((*21*) (*22*) (*23, 24*)).

To address this question, human CTLs were loaded with WGA-AF488 followed by staining with Vybrant™ DiI and Actin SPY650, seeded on coated ibidi chambers and inspected using TIRFM. As shown in **Figure 2a** and **Supplementary Movie 7** and **8**, thinning out of the actin cytoskeleton network overlapped with the hot-spots of SMAP release within areas of reduced membrane staining. Cortical actin hypodense areas, and corresponding membrane reduced staining, exhibited various morphology and dimensions in individual cells, with some cells showing central cSMAC-like secretion patterns (**Figure 2a**, Cell-1) and others exhibiting multiple secretion hot-spots (**Figure 2a**, Cell-2). These features were not observed in unstimulated cells (**Supplementary Movie 9**).

**Figure 2.**
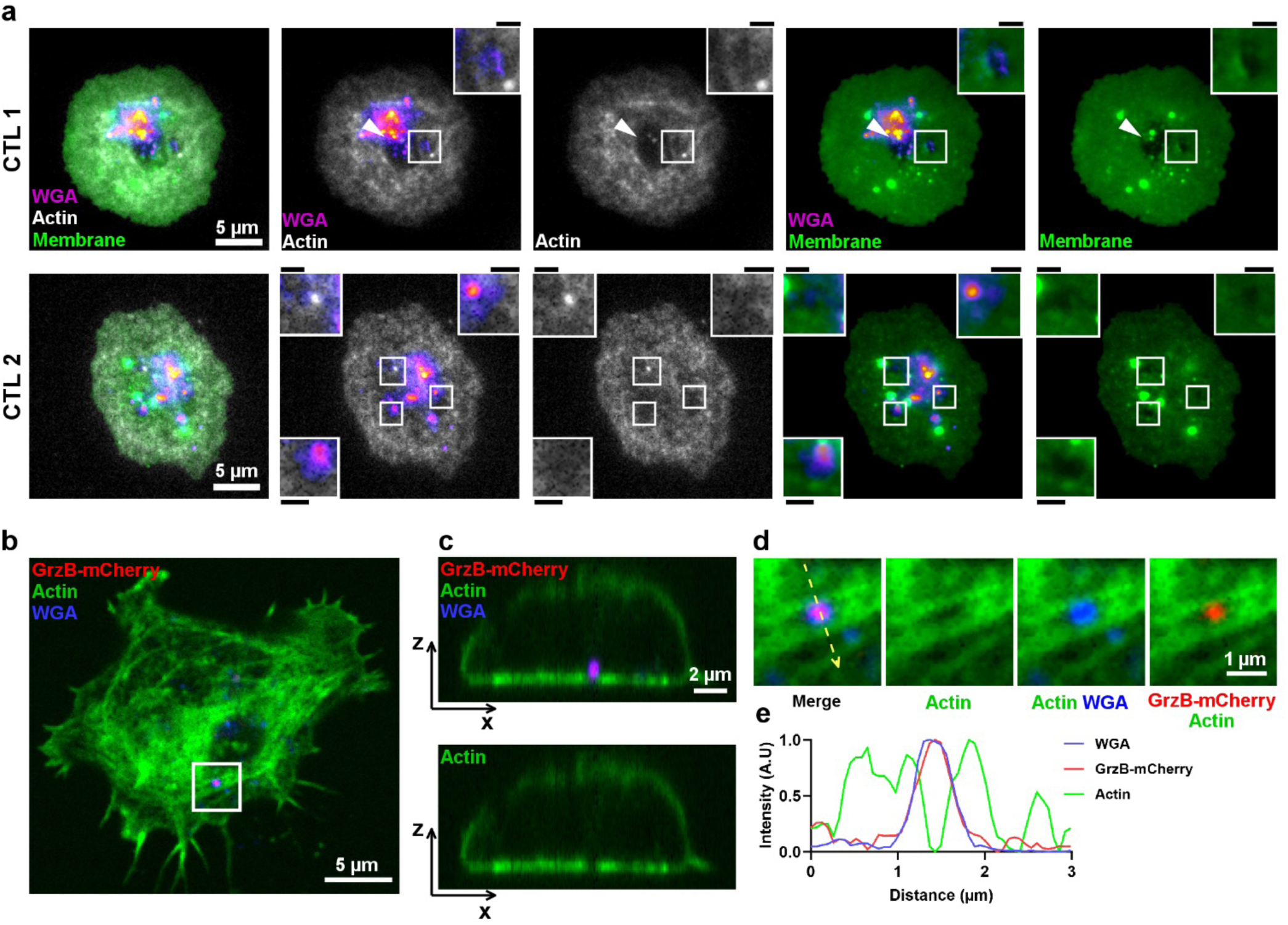
Secreted SMAPs percolate through the CTL actin cytoskeleton cortical network. **(a)** Live TIRF imaging of CTLs seeded on CD3/ICAM-1 coated ibidi µ-slide chambers at 37°C/5%CO_2_. CTLs were loaded with WGA-AF488 (pseudocolor) and stained with Vybrant^TM^ DiI (green) and Actin SPY650 (white). Upper and lower panels show two representative cells exhibiting different patterns of cortical actin depletion and corresponding membrane reduced staining. cSMAC-like secretion areas in Cell-1 (upper panels); multiple secretion hot-spots in Cell-2 (lower panels). Images are from **Supplementary Movie 7 and 8**. Data are from one representative experiment out 2. **(b)** Shown is a structured illumination 3D spinning disk image of a representative CTL transfected with RNA coding for GrzB-mCherry (red), pre-loaded with WGA-AF647 (blue) and stained with CellMask^TM^ Green Actin (green). The white square marks the area magnified in (d). **(c)** x/z projection depicting a WGA+/GrzB+ SMAP particle crossing an area of actin clearance. **(d)** Magnification of the region delimited by the white square in (b) depicting a SMAP particle navigating through the actin cytoskeleton network. **(e)** Plot profile of the normalized intensity (arbitrary unit) of WGA, GrzB and actin along the yellow dashed arrows in (d). Data are from one representative experiment out 2.

We next investigated whether granules carrying WGA^+^ particles also contained other SMAP components such as granzyme B (GrzB). To this end, CTLs were transfected with RNA coding for GrzB-mCherry and loaded with WGA-AF647 and CellMask™ Green Actin. As shown in **Supplementary Movie 10,** cells were successfully transfected with GrzB-mCherry RNA and exhibited a considerable fraction of GrzB^+^WGA^+^ SMAPs containing granules among the total vesicles of each cell.

When the transfected cells were stimulated as described in Fig 1, then fixed and imaged using spinning disk Live LSR imaging, (an approach equivalent to structured illumination 3D imaging, (*25*), GATACA Systems) WGA^+^GrzB^+^ SMAPs could be beautifully captured in their navigation through the actin cytoskeleton network (**Figure 2b-e** and **Supplementary Fig. 2**). Under these conditions, the areas of reduced density of actin cytoskeleton appeared narrower when compared to those observed in TIRF images on live CTL. Differences might be due to the higher resolution of structured illumination imaging in fixed cells that might allow detection of thin actin cytoskeleton meshwork within the actin hypodense areas ((*22*) (*23, 24*)).

All in all, the above results show that shortly after TCR triggering, human CTLs release SMAPs within areas of enhanced membrane turnover and matching areas of cortical actin cytoskeleton network re-modeling.

### SMAPs re-polarize towards lytic synapse and penetrate into target cells within minutes after cell-cell contact

We next investigated the time/space parameters of SMAP secretion in human CTLs stimulated by HLA-matched target cells (either JY, an EBV-transformed B cell line or MDA-MB-231, a triple negative breast cancer cell line) pulsed or not with an antigenic peptide, a condition that mimics physiological CTL activation via TCR engagement.

In an initial approach, we used CLEM to visualize SMAP release within the synaptic cleft. To this end, antigen specific CTLs previously transfected with GrzB-mCherry and loaded with WGA-AF488 were conjugated for 15 minutes with JY cells and processed for CLEM (**Figure 3a)**. This approach allowed us to detect in the synaptic cleft and within minutes after TCR engagement, numerous non-membrane bound electron dense particles with a size compatible with SMAPs (**Figure 3b and c**, displaying higher magnification of the region delimited by the white rectangle in 3a). These particles were found to be positive for WGA, GrzB or both (**Figure 3b** cyan, magenta and violet arrows respectively). SMAP release was not observed in conjugates in which target cells did not display antigenic determinants (**Figure 3d**).

**Figure 3.**
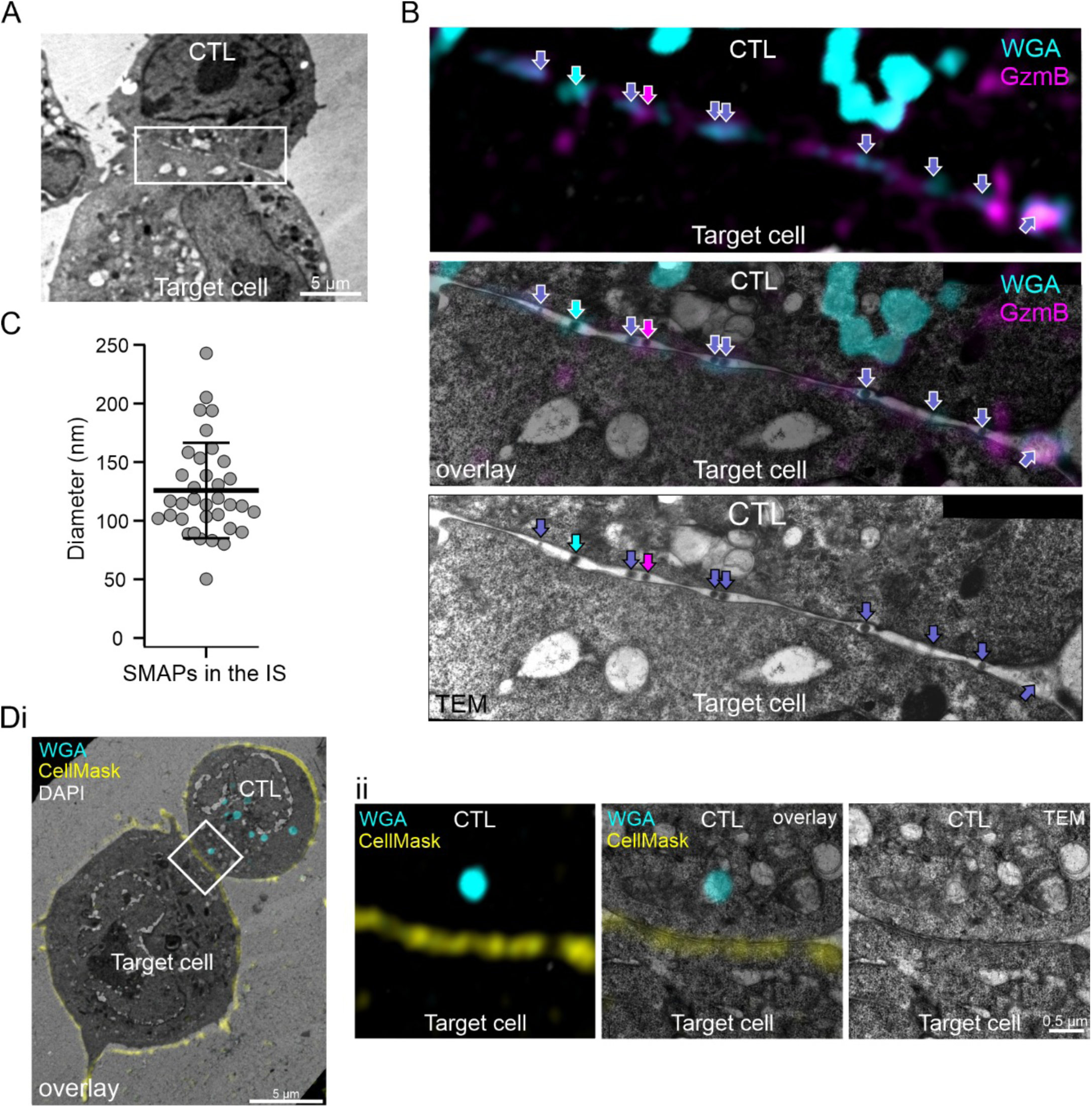
WGA^+^, GzmB^+^ and double positive SMAPs appear in the synaptic cleft within minutes after TCR engagement. **(a)** Electron micrograph of a representative CTL/target cell conjugate fixed and imaged after 15 min of contact. The white rectangle marks the area magnified in (b). **(b)** Correlative light electron microscopy (CLEM) analysis of the magnified immunological synapse of the CTL/target cell conjugate shown in (a) (white rectangle). The CTL was transfected with RNA coding for GzmB-mCherry (561 nm, magenta) and pre-incubated with WGA-AF488 (cyan). From top to bottom, the structured illumination microscopy (SIM) image, the CLEM image, and the electron micrograph are shown. The arrows mark WGA^+^ and GzmB^+^ SMAPs with the corresponding color code. GzmB^+^/WGA^+^ particles are marked with a purple arrow. **(c)** Diameter analysis of WGA^+^, GzmB^+^, and WGA^+^/GzmB^+^ SMAPs present in the lytic synapse (LS). Data are given as mean ± SD; N_LS_=6, n_SMAPs_=35. **(di)** CLEM analysis of a representative non-pulsed CTL/target cell conjugate (left panel). CTL was preloaded with WGA-AF488 (cyan). The white rectangle marks the region of interest magnified in (dii). **(dii)** From left to right: SIM image, CLEM overlay and electron micrograph of the magnified contact zone between CTL/target cell shown in (di). CellMask^TM^ Deep Red (yellow) and DAPI (white) signals correlate to the plasma membrane and the nucleus, respectively.

While CLEM approach allowed us to provide ultrastructural evidence of SMAP release during cognate interaction of CTLs with target cells, it did not allow us to capture the full 3D synaptic structure.

To overcome this limitation, we studied SMAP synaptic release using spinning disk Live LSR microscopy, an approach equivalent to structured illumination 3D imaging ((*25*), GATACA Systems). CTL were pre-loaded with WGA-AF647, conjugated with JY target cells, fixed at different time points, stained with anti-GrzB, anti-CD107a and phalloidin.

As shown in **Figure 4a**, CTL interacting with unpulsed target cells did not polarize their SMAP containing CD107a^+^ lytic granules towards the cell-cell contact sites (**Figure 4a left panel)**. Under these conditions a clear heterogeneity of cytotoxic content in lytic granules was observed in individual CTLs (**Figure 4a left and central panels and Supplementary Movie 11**). WGA^+^ and GrzB^+^ particles were exclusively detected within CD107^+^ granules which were heterogeneously distributed in each individual CTL. Some granules were GrzB^+^ only (possibly corresponding to single core granules (SCG) that release soluble GrzB), others were WGA^+^ only and some were double positive for GrzB^+^ and WGA^+^, most likely containing SMAPs (**Figure 4a right panels** and **Supplementary Movie 12,** see also **Figure 4b right panel**).

**Figure 4.**
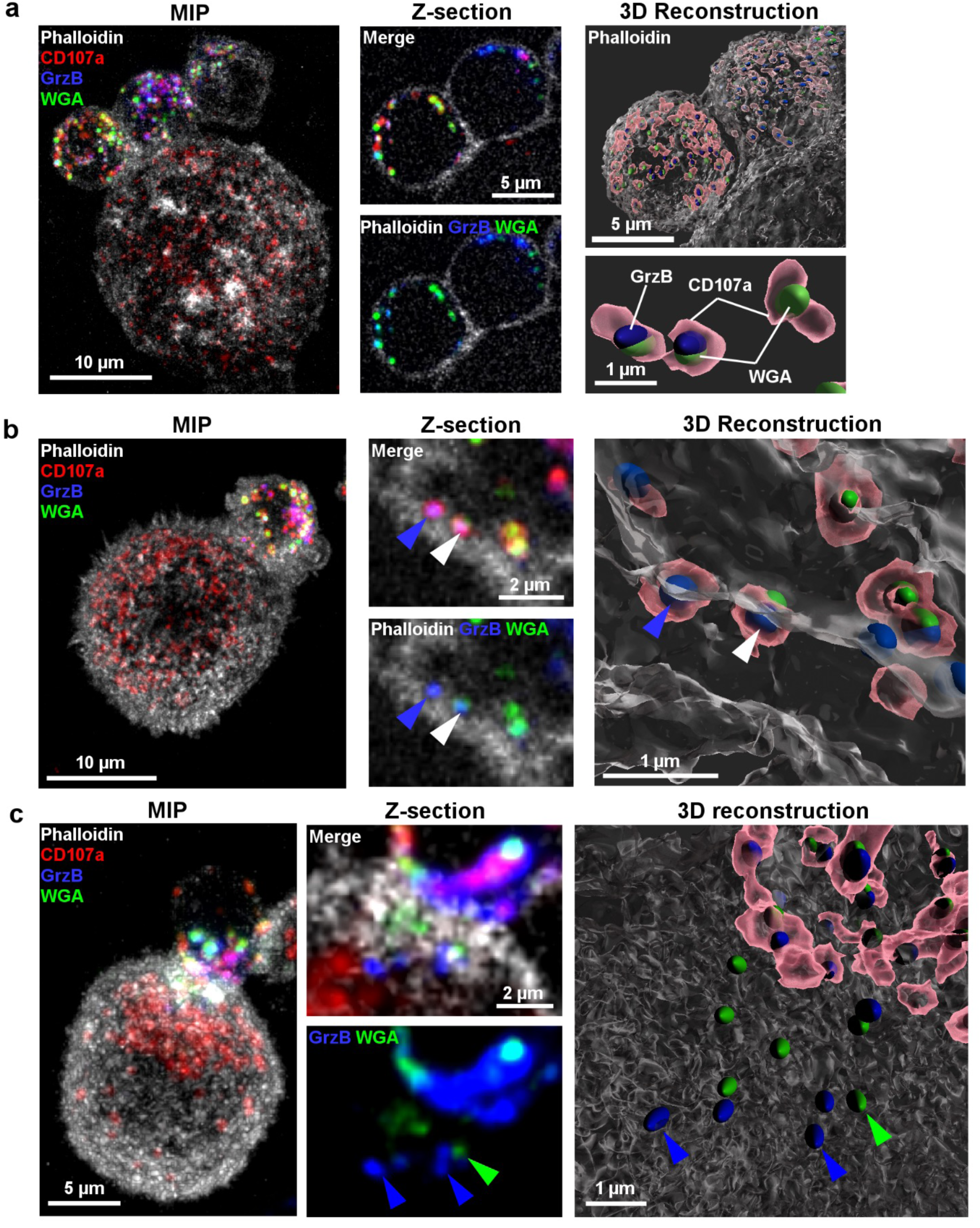
SMAPs re-polarize towards lytic synapse and penetrate into target cells within minutes after cell-cell contact. **(a)** Left panel: Maximum Intensity Projection (MIP) of a representative conjugate formed between a CTL and a target cell (JY) displaying no antigenic peptide. CTLs were previously loaded with WGA-AF647 (green), fixed and stained with phalloidin (white) and antibodies directed against CD107a (red) and GrzB (blue); middle panels: z-sections highlighting the heterogeneity of the lytic granule content of individual clonal CTLs; right panels: 3D image reconstruction depicting representative distribution in CTLs of CD107a^+^ vesicles (red) containing GrzB (blue) and WGA (green) particles. Images are from **Supplementary Movie 11** and **12**. **(b)** Left panel: MIP of a representative conjugate formed between a CTL and a peptide-pulsed target cells interacting for 2 minutes. Conjugates were fixed and stained as in (a). Middle panel: Magnification of a z-section of the contact area between CTL and target cell. WGA^+^, GrzB^+^ or GrzB^+^WGA^+^ particles are detected in the cortical actin network of the CTL (blue and white arrows); right panel: 3D reconstruction showing SMAP particles still contained within CD107a+ vesicles passing through CTL-actin cortical network. Images are from **Supplementary Movie 13.** **(c)** Left panel: MIP of a representative conjugate formed between a CTL and a peptide-pulsed target cells interacting for 30 minutes. Conjugates were fixed and stained as in (a). Middle panel: Magnification of a z-section of the contact area. WGA^+^, GrzB^+^ or GrzB^+^/WGA^+^ particles were detected passing through the synaptic cleft and inside of the target cells (blue and green arrows). Images are from **Supplementary Movie 14**); right panel: 3D reconstruction depicting SMAPs inside the target cell (blue and green arrows). Data are from one representative experiment out of 5.

In contrast, during cognate interaction of CTL with target cells, SMAP containing granules polarized towards lytic synapses in a time dependent fashion (**Figure 4b-c**). Remarkably, as early as 2 minutes after conjugation, WGA^+^, GrzB^+^ and WGA^+^GrzB^+^ granules could be detected within the cortical actin network of CTLs in the process of being transferred from CTLs to target cells (**Figure 4b central and right panels** and **Supplementary Movie 13**). At later time points (30 minutes), we SMAPs were detected passing through the synaptic cleft and internalized by target cells (**Figure 4c central and right panels** and **Supplementary Movie 14**).

To quantitatively asses the rapid re-polarization of SMAPs containing lytic granules and the penetration of single or double positive particles into target cells, we analyzed a statistically relevant number of CTL/target cell conjugates in 3D using Imaris software as depicted in **Supplementary Figure 3**. As shown in **Supplementary Figure 4a**, a clear time-dependent polarization of SMAPs was observed as measured by decreasing distance from cortical actin cytoskeleton over time. Negative values in the panels indicate SMAPs detected within the cortical actin network of the contiguous cells.

Similar results were obtained when CTLs were conjugated with the HLA-matched triple negative breast cancer cells MDA-MB-231 pulsed or not with the antigenic peptide (**Supplementary Figure 4b**).

The above results show that SMAPs repolarize towards lytic synapse rapidly after TCR engagement and are internalized by target cells within the initial 30 minutes after cell-cell encounter.

### SMAP enter target cell endosomal compartment

We next investigated the fate of SMAPs once inside target cells. It is well established that peforin-induced membrane damage triggers an active endo/exocytic process to repair the cell membrane ((*12, 13*)). We therefore hypothesized that SMAPs might gain access to target cells cytosol through this repair associated endocytic pathway. To verify this hypothesis, we focused on MDA-MB-231 cells which exhibit very abundant and large vesicles of the endocytic compartment as detected by staining for markers of early and late endosomes. Human CTLs were loaded with WGA-AF647 and conjugated with MDA-MB-231 cells for 1 hour. Cells were fixed, permeabilized and stained with phalloidin and antibodies directed against EEA1 or CD107a to detect early endosomes or lysosomes/late-endosomal vesicle (LLE), respectively. As shown in **Figure 5a** and **c** this analysis revealed, that a fraction of WGA^+^ particles was detected in the CD107a^+^ target cell compartment upon interaction with cognate CTLs. Staining for EEA1 provided complementary results (**Figure 5a** and **c**). To better visualize WGA^+^ particles within early and late endosome vesicles we performed 3D reconstruction of the images using Imaris software (**Figure 5b**). To provide quantitative 3D evaluation of WGA^+^ particle inside of vesicular compartments we measured the overlapping of WGA^+^ particles with EAA1 or CD107a (**Figure 5d-g**). **Figure 5e** and **f** show examples of the measured overlapping between WGA and markers of endosomal compartment as detected in Imaris 3D reconstructions.

**Figure 5.**
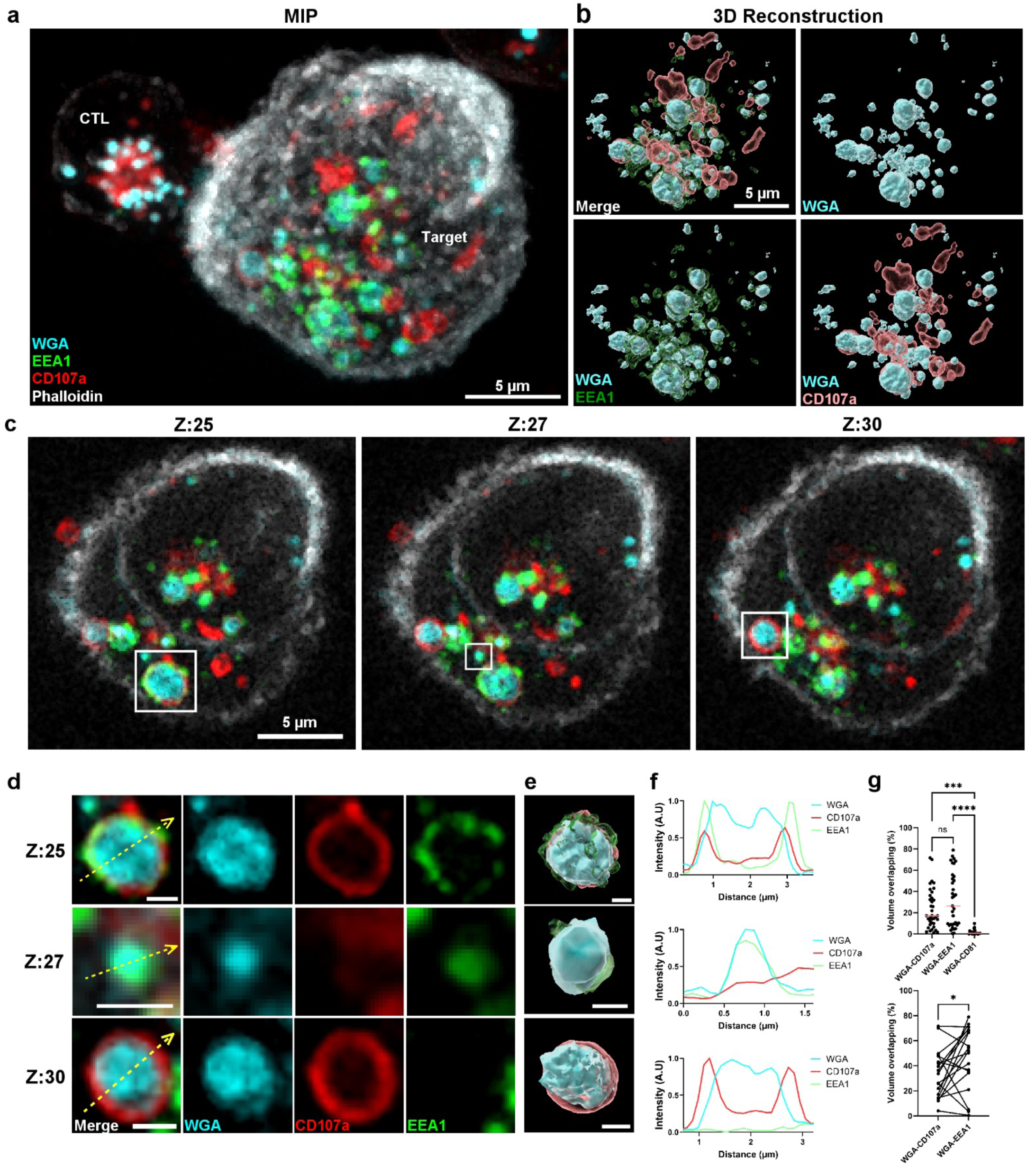
SMAPs are taken up into the endosomal compartment of target cells. **(a)** MIP of a representative conjugate formed for 1h, between a target cell (MDA-MB-321) and a previously WGA-AF647 (cyan) loaded CTL. Conjugates were fixed, permeabilized and stained with phalloidin (white) and antibodies against EEA1 (green) and CD107a (red). Data are from one representative experiment out of 4 for CD107a, 3 for EEA1 and 2 for both markers stained simultaneously. **(b)** 3D reconstruction showing the overlaps of internalized WGA^+^ particles with EEA1/CD107a (merge, upper left panel), EEA1 (green) or CD107a (red) (lower left and right panels respectively) and the WGA^+^ particles alone (cyan, upper right panel). **(c)** Tree individual z-sections of the MIP in (a) illustrating variable WGA overlapping with individual early and late endosomal vesicles. The white squares mark the areas magnified in (d). **(d)** Magnification of the regions delimited by the white squares in (c) showing for each z-section, WGA (cyan), CD107a (red) and EEA1 (green) staining separately. **(e)** 3D reconstruction of the individual compartments. **(f)** Plot profile of the normalized intensity (arbitrary unit) of WGA, EEA1and CD107a along the yellow dashed arrows in images (d). **(g)** Percentage of overlapping volume of WGA with the volume of CD107a, EEA1 and CD81 inside the target cell (upper panel). Correlation between the percentage of overlapping volume of WGA with the volume of CD107a and the WGA overlapping volume with EEA1 in individual cells (lower panel). Measurements in (g) are from 2 independent experiments. Upper panel N=37 for WGA-CD107a, N=35 for WGA-EEA1 and N=19 for WGA-CD81. Lower panel N=19. Statistical analyses were performed using GraphPad Prism 10.1.2. Upper panel, one-way ANOVA was utilized to assess statistical differences. \**p*<0.05, ** *p*<0.01, *** *p*<0.001, \*\*\*\**p*<0.0001, ns = not significant. Lower panel, a parametric paired Student’s t-test was utilized to assess statistical differences. \**p*<0.05.

The 3D analysis revealed that in each target cell about 30% of WGA^+^ particles overlapped with EEA1 staining while less than 20% were found in association with CD107a^+^ vesicles with a large variability from one target cell to the other (**Figure 5g** upper panel). Interestingly, in individual target cells an inverse correlation between the localization of WGA into CD107a^+^ vesicles and EEA1^+^ vesicles was observed, indicating that in each cell SMAPs are differently distributed within the two endosomal compartments (**Figure 5g** lower panel). Conversely WGA^+^ particles were not found in association with the multivesicular body/exosome marker CD81 suggesting that this compartment is not involved in SMAP trafficking in target cells (**Figure 5g** upper panel).

Globally, the above measurements revealed that in snapshot images collected 1 hour after conjugate formation a large fraction of internalized SMAPs is found within the early/late endosomal compartment of target cells as detected by EEA1 and CD107a staining.

To investigate the time-kinetics of this phenomenon WGA-loaded CTLs were conjugated for different times with peptide pulsed MDA-MB-231 cells. Conjugates were fixed, permeabilized and stained with phalloidin and antibodies directed against GrzB and CD107a. As shown in **Figure 6a** CTL rapidly polarized their SMAPs containing lytic granules towards cognate target cells, which in turn deployed their LLE vesicle based synaptic defense, in agreement with our previous findings ((*12, 13*)). 3D visualization of the images revealed a time-dependent transit of WGA^+^GrzB^+^ particles from the CTL CD107a^+^ compartment (red) into the target cell CD107a^+^ compartment (blue), with detectable particle transfer at 30 minutes.

**Figure 6.**
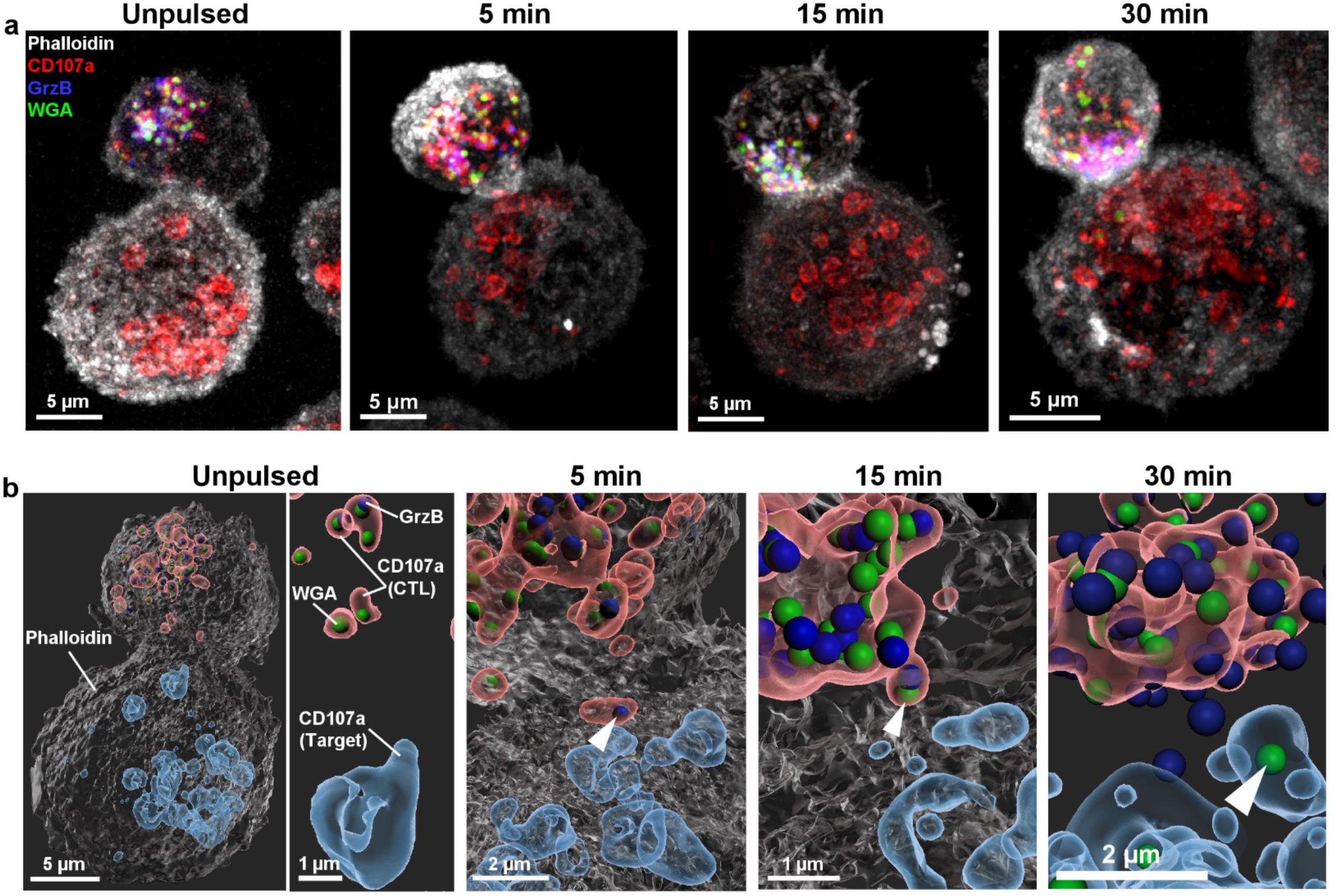
Target cell endocytic pathway represents a time dependent gateway for SMAP internalization. **(a)** MIP of conjugates formed between CTL and MDA-MB-231 cells either unpulsed (left) or pulsed with the specific antigenic peptide. Images are from different time points. Conjugates were fixed and stained as in Figure 4. **(b)** 3D visualization of the time-dependent transit of WGA^+^(green) and GrzB^+^ (blue) particles from the CTL-CD107a^+^ compartment (red) into the target cell-CD107a^+^ compartment (sky blue). White arrows indicate WGA^+^and GrzB^+^ particles within CD107a^+^ vesicles in target cells. Data are from one representative experiment out of 2.

Taken together the above results indicate that the target cell endosomal compartment represents a gateway for SMAP internalization.

### Intercellular SMAP transfer via stenocytosis

Together, the above observations showed that SMAPs rapidly exit CTLs trough localized regions of actin depletion areas and are subsequently taken up by target cells at the lytic synapse. They raise the question of by which mechanism SMAPs penetrate into target cells. To address this question, we focused on the organization of target cell actin cytoskeleton, as it has been previously reported to act as a barrier for the penetration of lytic molecules during NK cell mediated cytotoxicity ((*26*)).

We therefore performed careful inspection of our CTL target cell conjugates at early and intermediate time points (2 to15 minutes) during which SMAP transfer is most evident. This approach allowed to capture “narrow passages” in the actin cytoskeleton through which particles appeared to move from one cell to the other (**Figure 7a-c** and **Supplementary Movie 15)**. Interestingly, these narrow passages appeared to traverse the entire thickness of the cortical actin cytoskeleton of the two cells in contact, suggesting that coordinated dynamic remodelling of actin at the synaptic cleft may transiently open conduits facilitating SMAP passage between the two cells.

**Figure 7.**
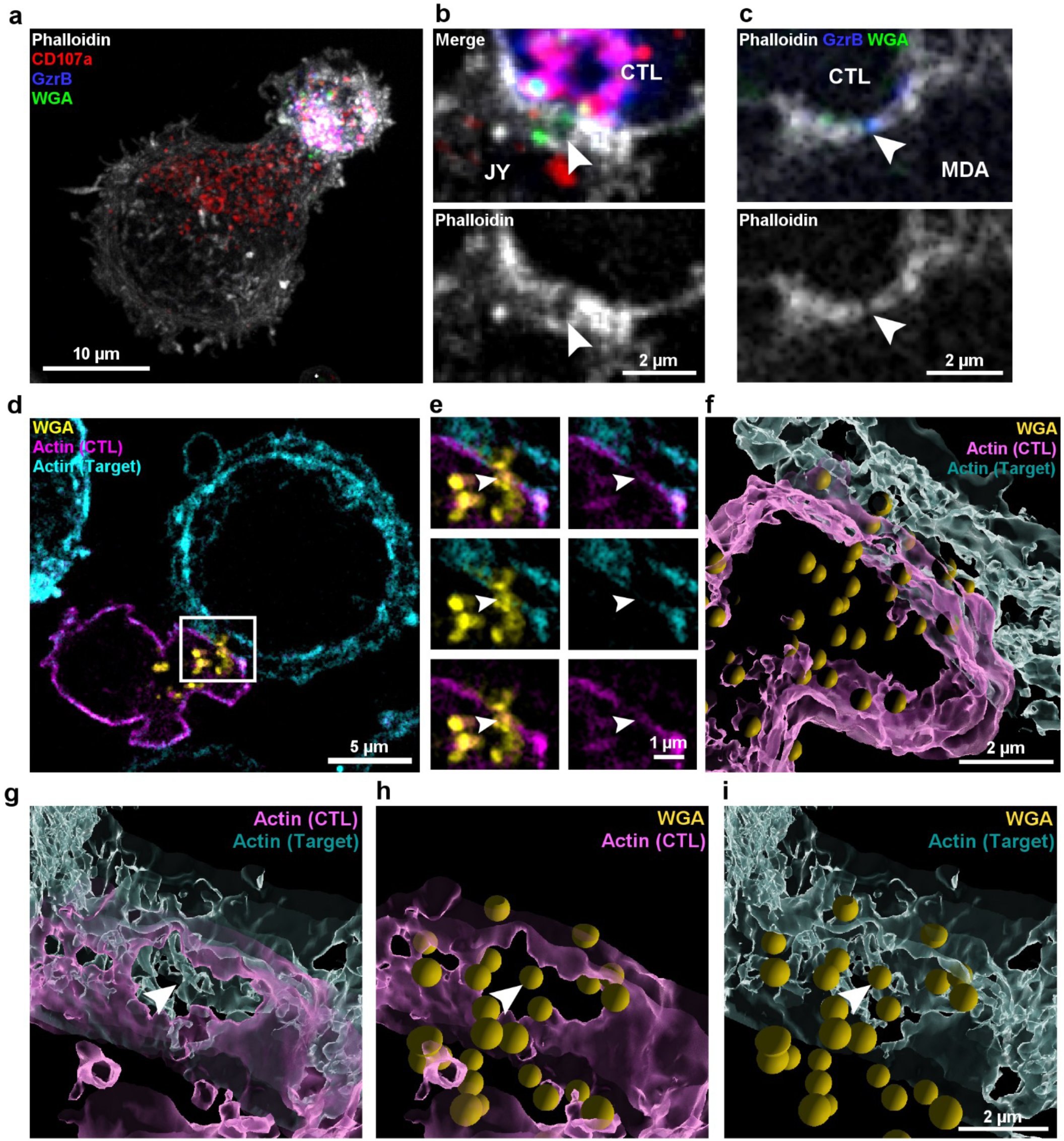
Intercellular SMAP transfer via *stenocytosis*. **(a)** Left panel: MIP of a representative conjugate formed between a CTL and a peptide-pulsed target cell (JY) interacting for 15 minutes. Conjugates were fixed and stained as in Figure 4. Images are from **Supplementary Movie 15.** **(b)** Magnification of a z-section of the contact area of the same conjugate shown in (a). SMAPs are detected passing through cortical actin interstices from CTL to target cell (white arrows). **(c)** Magnification of a z-section of the contact area between a CTL and a MDA-MB-231 conjugated for 2 minutes. SMAPs are detected passing through cortical actin interstices from CTL to target cell (white arrows). Data are from one representative experiment out of 5 for 15 minutes and 3 for 2 minutes. **(d)** A representative conjugate formed between a CTL (previously loaded with CellMask^TM^ Orange Actin, magenta) and WGA-AF647 (yellow) and a target cells (JY) loaded with CellMask^TM^Green Actin (cyan). The white rectangle marks the area magnified in e-i. **(e)** Magnification of the region delimited by the white square in (d). White arrows point the actin clearance areas in the CTL and in in the target cell. **(f)** 3D visualization of the immunological synapse formed between the two cells (CTL, magenta; target cell, cyan), depicting the passage of WGA^+^ particles (yellow) via *stenocytosis*; **(g)** CTL (magenta) and a target cell (cyan) actin cytoskeleton. **(h)** CTL (magenta) actin cytoskeleton and WGA^+^ particles (yellow). **(i)** Target cell (cyan) actin cytoskeleton and WGA^+^ particles (yellow). Data are from one representative experiment out 3.

The to define the frequency of these narrow cortical actin depletion areas, inspection across the acquired z-stacks of CTL/target cell (JY and MDA-MB-231) conjugates was performed separately by two individuals. This analysis showed that out of 103 scored CTL/JY conjugates, 34 exhibited (33%) detectable actin depletion areas. For 108 scored CTL/MDA-MB-231 conjugates 32 (29%) were positive for the actin cell-cell “narrow passages” To better characterize the phenomenon of SMAP intercellular transfer, we differentially labeled CTL and target cell actin cytoskeleton with either Cell Mask™ Orange actin (violet) or Cell Mask™ Green actin (cyan) prior to their conjugation. This dual color labeling enabled clear discrimination of the cortical actin cytoskeleton of the two adjacent cells. CTL were previously loaded with WGA-AF647(yellow) to detects.

This experimental approach showed that in CTL/target cell conjugates SMAPs moved from one cell to the other via corresponding narrow passages formed in the cortical actin cytoskeleton of both cells (**Figure 7d and Supplementary Figure 5a and b** in which three typical conjugates are depicted). Synaptic blow ups allowed to better appreciate SMAPs cell-cell transfer within actin cytoskeleton hypodense areas (**Figure 7e and Supplementary Figure 5a and b**, white arrows). To have a 3D view of the matching hypodense areas in the cortical actin cytoskeleton of the two cells we reconstructed the synaptic areas using Imaris. This analysis allowed us to illustrate SMAPs transfer trough matching openings formed in the cortical actin cytoskeleton of the two adjacent cells (**Figure 7f-i**).

Collectively, these findings demonstrate a dynamic secretion mechanism at the CTL/target cell interfaces, driven by the rapid transport of SMAPs within CD107a⁺ granules in CTLs, their passage through localized narrow regions of cortical actin depletion in both cells and subsequent uptake into the target cell’s endosomal compartment. We have termed this previously unrecognized mode of intercellular transfer of supramolecular clusters as ***stenocytosis*** (see **Supplementary Figure 6**).

## Discussion

The interaction between CTLs and target cells has been traditionally seen as a pray/predator scenario in which packs of highly efficient killer cells attack easy preys and eliminate them. Research over the last three decades in the field of tumor immunology inspired a substantial revision of this paradigm by showing that both killer and tumor target cell populations are highly heterogeneous ((*10, 27–29*)). Consequently, during the interaction between CTLs and tumor target cells, the efficacy of cytotoxicity results from the lytic potential of individual killer cells, the resistance of individual target cells to cytotoxic attack and the number and frequency of encounters between CTLs and target cells. ((*2*) (*30*)).

Nowadays, CTL/tumor target cell interaction is seen as a highly complex multistep process that is essential to unravel in order to design immunotherapy strategies able to shift the balance between CTL efficacy and tumor target cell resistance at the lytic synapse in the favor of CTLs ((*31*)). The discovery of SMAPs has raised interest in the potentially groundbreaking immunotherapeutic power of these autonomous killing entities. Yet the relevance of SMAPs in synaptic lethal hit delivery and their contribution to overcome synaptic tumor cell resistance to CTL attack is presently unknown.

To shed new light on SMAP synaptic function we investigated, in antigen specific human CTLs, SMAP intracellular dynamics and release together with their subsequent uptake into target cells. We show that SMAPs can serve as directionally secreted synaptic weapons. They are detectable in the synaptic cleft within minutes after CTL encounter with cognate target cells and are rapidly taken up into the target cell endocytic compartment via a previously unidentified process of *stenocytosis,* based on coordinated formation of hypodense areas in opposing CTL and target cell cortical actin cytoskeleton (**Supplementary Fig. 6**).

In the initial part of our study, we focused on the kinetics of SMAP release by human CTLs triggered by immobilized anti-CD3/ICAM-1, a surrogate stimulus mimicking CTL interaction with cognate target cells and TCR-mediated antigen recognition. We employed TIRFM to monitor, with high time/space resolution, the kinetics of SMAPs release by individual CTLs.

A first finding provided by this approach was that human CTL possesses a large fraction of preformed SMAPs, which are promptly released within a few minutes after TCR triggering. This observation was confirmed by the detection of swift SMAP containing granule re-polarization and release upon CTL interaction with cognate target cells (**Figure 3** and **4** and **Supplementary Figure 4**). SMAPs have been initially described as weapons in the CTL tactical arsenal acting in between the two previously reported time scales of killing: the release of soluble cytotoxic proteins from dense core granules into the immunological synapse (seconds/minutes) and the FasL-mediated apoptosis (hours/days) ((*3*))(*32*)). Our present findings extend this concept by showing that SMAP release by human CTLs can be very rapid, highlighting their potential as an early synaptic weapon. A second result provided by the study of CTL interacting with immobilized anti-CD3/ICAM-1 is that SMAPs release happens within areas of intense secretion activity that match areas of cortical actin cytoskeleton depletion. This observation is in line with previously reported data showing that, in mouse CTLs, a PIP_2_-dependent synaptic actin depletion/recovery is mechanistically linked to the initiation and termination of lytic granule secretion ((*21, 33*)).

It is interesting to note that in our study each CTL presents different patterns of SMAP secretion: with some cells exhibiting cSMAP-like central areas of secretion and others presenting multiple small areas of hypodense cortical actin. This observation suggests that, in CTLs SMAP secretion can stochastically occur in multiple zones of the activation contact after TCR triggering.

As such, our results are compatible with a model in which, in human CTL, cortical actin cytoskeleton might serve as a physical barrier to prevent docking and fusion of secretory vesicles, including the SMAPs carrying MCGs, at the plasma membrane ((*21*) (*20*).

Nevertheless, it should be noted that our TIRF-based imaging of live cells, while exhibiting high time resolution, might lack the space resolution and sensitivity to detect the cortical actin network in the actin hypodense areas. Our observations are therefore compatible with the possibility that cortical actin cytoskeleton might not be completely removed from synaptic hypodense areas. Conversely, the remodeling of synaptic cortical actin meshwork might play a key role in lytic granule docking and secretion as previously reported in NK cells ((*22, 23, 34*) (*24*)). This hypothesis is in line with our observation that in fixed CTLs interacting with immobilized stimuli (**Figure 2b-d**) and in fixed CTL/target cell conjugates (**Figure 7**), imaged by structured illumination microscopy, cortical actin presented relatively narrow areas of actin depletion allowing transfer of SMAPs.

A third result obtained when monitoring human CTL stimulation with immobilized ligands is that, using rapid 3D spinning-disk microscopy, we could image instant SMAP-containing vesicles docking and fusion with plasma membrane via a secretion mode reminiscent of the Ω-profiles described in exo/endocytosis of neuro-endocrine cells ((*18*) (*19*) (*20*)). It is tempting to speculate that a secretory mechanism consisting of vesicle shrink fusion with formation of Ω-profiles allowing the release of quanta of bioactive material may underlie the discrete release of bioactive substances such as hormones, neurotransmitters, and immune mediators.

The study of SMAP release in cognate CTL /target cell conjugates provided complementary results and unveiled previously unidentified features of cell-cell communication at the lytic synapse. CLEM imaging allowed the first description of SMAPs within human CTL target cell synaptic clefts. It is tempting to speculate that these hundred nanometers-sized particles present at the synapse might contribute to the plasma membrane spacing and interdigitation observed only in antigen specific conjugates and not in unpulsed conjugates in which, cell-cell contacts, were on the contrary linear and tight. The presence of these compact supramolecular particles at the synapse might locally interfere with TCR/pMHC engagement at the synapse by creating nano-areas of intermittent TCR triggering within synapses, a mechanism that would contribute to pace the sustained signaling and secretory process.

The time kinetics of re-polarization of SMAP containing CD107a^+^ vesicles towards the lytic synapse and of SMAP passage into target cells, as detected by structure illumination microscopy, shows a prompt response initiated within minutes. This time frame is compatible with the time required for the re-polarization of the CTL microtubule organizing center (MTOC) towards the synapse area as reported by others and us ((*8, 35, 36*)). It is therefore likely that SMAPs carried by MCGs might not be released within seconds after TCR engagement as we previously described for soluble perforin resulting in membrane pore formation, [Ca^2+^]_i_ waves in target cells and initiation of Ca ^2+^ depended target cell reparative membrane turnover ((*13*)). On the contrary SMAPs carrying granules appear to reach the synaptic area only minutes after soluble perforin has started to work. Their polarization and secretion might be therefore more tightly bound to MTOC repolarization.

All in all, our results position SMAP release right after the ultra-rapid perforin release ((*8*)) with SMAPs being part of a second lytic wave that re-enforces the first wave of attack mediated by soluble cytotoxic components released by CTLs.

It is interesting to note that target cells polarize and secrete their LLE vesicles at the lytic synapse in the context of a Ca^2+^ dependent membrane reparation mechanism. We initially described this phenomenon in melanoma cells ((*12, 13*)) however, many target cells exhibit this rapid response as shown in our current images (**Figure 6a** and **7a**). We hypothesize that the active exo-endocytic process occurring on the target cell side of the lytic synapse might favor SMAP capture by the target cells. Accordingly, we find that SMAPs enter target cells via their endosomal compartment (**Figure 5**).

Having observed that SMAPs are released into the synaptic cleft within minutes after CTL/target cell contact and that they enter the target cell endosomal compartment we asked how SMAPs might traverse the target cell cortical actin cytoskeleton that has been reported to act as a defensive shield protecting target cells from NK cell attack ((*26*)). Surprisingly our findings show that, target cell cortical actin is permeable to SMAPs and exhibit narrow areas of reduced actin density in correspondence with SMAPs release (**Figure 7**). This observation is puzzling and raises the question of which molecular mechanism might be responsible of the observed cytoskeleton response in target cell SMAP *stenocytosis* from one cell to the other. Further research is required to identify the precise signals leading to target cell actin network density reduction following CTL attack. It is tempting to speculate that the active Ca^2+^-dependent secretory/endocytic process occurring in cognate CTL/target cells might cause the formation of actin cytoskeleton hypodense areas mirroring those formed in CTLs to allow target cell endosomal vesicles docking, fusion and retrieval via mechanisms similar to those described in other secretory cells.

Our results extend those previously obtained on the actin-mediated defense of target cells (((*26*)). They suggest that while in certain tumor cell types and in individual cells belonging to a cancer cell population actin cytoskeleton polymerization can act as a shield to limit soluble perforin/granzyme-mediated target cell killing, in other cells SMAPs can find their way to penetrate cortical cytoskeleton to elicit cell death. Finally, they imply that actin cytoskeleton-mediated defense and LLE secretion defense mechanisms might be somehow in competition and be differently used by individual target cells facing CTL.

More in general our research adds to the notion that the cortical actin cytoskeleton plays a key role in the dynamics of CTL/target cell interaction and in various aspects of T cell activation by controlling several key functions including cell-cell adhesion, molecular segregation and mechanical force generation ((*37, 38*) (*39*)).

Collectively our data identify SMAPs as second line cytotoxic weapons of CTLs readily deployable against target cells in order to re-enforce cytotoxic outcome. This observation raised the question of whether SMAP release at the synapse may differently contribute to the cell death process of target cells when compared to the ultra-rapid lethal hit mediated by soluble lytic molecules.

An answer to this question comes from a recent study from our lab in which we show that perforin-mediated pore formation in target cell membrane induces an early PCD that is pyroptotic and pro-inflammatory. Conversely, isolated SMAPs trigger a PCD that is mostly apoptotic (https://doi.org/10.1101/2024.05.24.595698 and our unpublished results). The observation that perforin itself and SMAPs released at the lytic synapse in a rapid sequence can autonomously engage different PCD pathways highlights the complexity and versatility of the CTL lytic arsenal when facing target cell resistances. We hypothesize that the two mechanism of cell death might synergize with SMAPs triggering apoptosis in target cells that failed to die by pyroptosis. As a consequence, while pyroptotic target cell death of some target cells might be pro-inflammatory and immunogenic, SMAP-induced apoptosis might serve as a source of tumor antigens for cross-priming by dendritic cells.

In conclusion, together with previous data, our study contributes to put forth a two-tier model of lytic granule-based lethal hit delivery by human CTLs. Within seconds after TCR triggering CTL release soluble lytic perforin and granzymes initiating a process of inflammatory cell death and at the same time triggering reparative membrane turnover and LLE vesicle secretion in target cells as an attempt to resist the CTL attack. During the reparation process, a second rapid and abundant wave of lytic granules (DCG and MCG) is secreted at the synapse resulting in release of SMAPs, an additional synaptic weapon. SMAPs sneak via cortical cytoskeleton interstices behind the defense lines exploiting the endosomal vesicles deployed for reparation as a Troy horse to enter target cells. This process re-enforces killing efficiency and promotes apoptotic cell death.

## Acknowledgements

We thank the members of the European Research Council ATTACK consortium for their discussion and Dr. Roxana Khazen and Dr. Loïc Duprè for their helpful critiques of this manuscript. We thank the flow cytometry and microscopy core facilities of the INSERM UMR 1037 (CRCT). Parts of the figures (e.g. organelles, membranes) were drawn by using pictures from Servier Medical Art, which is licensed under a Creative Commons Attribution 3.0 Unported License. This research in this manuscript has received funding from the European Research Council (**ERC**) under the European Union’s Horizon 2020 Research and Innovation Programme (Grant agreement No. Syn-951329). The research was also supported by INSERM institutional funding, from the Fondation Toulouse Cancer Santé and from the Oncopole Claudius-Regaud. Funders had no role in the preparation of this manuscript.

## Author contributions

LD designed the research, performed experiments, analyzed results, and wrote the paper. CS, MPP, and SM performed experiments and analyzed data. BM discussed data and edited the paper. MLD and UB supervised research, discussed results and edited the paper. SM and SV supervised the research, designed the project, and wrote the paper.

## Data Availability

All original data generated for this study will be deposited in ZENODO repository server (https://zenodo.org/)with a unique identifier listed here upon acceptance of this article.

## Materials and Methods

### Cell Culture

Cell culture was performed as previously described. Human CD8^+^ T cells were purified from healthy donor blood samples using the RosetteSep Human CD8^+^ T Cell Enrichment Cocktail (StemCell Technologies). For cloning, HLA-A2-restricted CD8^+^ T cells specific for the NLVPMVATV peptide of the cytomegalovirus protein pp65 were single-cell sorted into 96-U-bottom plates using a BD FACSAria II cell sorter using tetramer staining. Cells were cultured in RPMI 1640 medium supplemented with 5% human AB serum (Inst. Biotechnologies J.BOY), 50µmol/L 2-mercaptoethanol, 10mM HEPES, 1X MEM NEAA (Gibco), 1X Sodium pyruvate (Sigma), 10µg/ml ciprofloxacine (AppliChem), 100 IU/ml human rIL-2 and 50 ng/ml human rIL-15 (Miltenyi). CD8^+^ T-cell clones were stimulated in complete RPMI/HS medium containing 1 μg/ml PHA with 1 x 10^6^ per ml 35 Gy irradiated allogeneic peripheral blood mononuclear cells from healthy donors (isolated on Ficoll Paque Gradient). Clones were restimulated every 2 weeks. Blood samples were collected and processed following standard ethical procedures after obtaining written informed consent from each donor and approval by the French Ministry of Research (transfer agreement AC-2020-3971). Approbation by the ethical department of the French Ministry of Research for the preparation and conservation of cell lines and clones starting from healthy donor human blood samples has been obtained (authorization no. DC-2021-4673). Two different target cells were used: JY, HLA-A2^+^ EBV-transformed B cells and MDA-MB 231, HLA-A2^+^ triple negative human breast cancer cell line. All target cells were cultured in RPMI 1640 GlutaMAX supplemented with 10% heat inactivated FCS (Gibco) and 50 µmol/L 2-mercaptoethanol, 10 mM HEPES, 1 X MEM NEAA (Gibco), 1 X Sodium pyruvate (Sigma), 10 µg/ml ciprofloxacine (AppliChem). All cell lines were screened biweekely for mycoplasma contamination using MycoAlert mycoplasma detection kit (Lonza).

### Total Internal Reflection Microscopy (TIRF)

Human CTL were loaded overnight with Wheat Germ Agglutinin Alexa Fluor 488 (1µg/mL) (Molecular Probes, Invitrogen) or WGA Alexa Fluor 647 (0,5µg/mL or 0,1µg/mL) washed and subsequently loaded with Vybrant™ Dil (Invitrogen) (1µM) and CellMask™ Green Actin Tracking Stain (5X) (Invitrogen™) or SPY620-actin (4X) (Spirochrome AG) for 30 minutes at 37°C and 5% CO_2_ in RPMI 5% FCS/HEPES. Cells were washed, resuspended in RPMI 1640 medium (1X) w/o pH Red supplemented with 10 mM L-Glutamine, 10 mM Hepes and 5% FCS and seeded into µ-Slide 15 Well 3D glass bottom slides (Ibidi) coated with poly-D-lysine (Sigma), human monoclonal anti-CD3 antibody (1 µg/mL, TR66, Enzo Life Sciences) and recombinant human ICAM-1/CD54 Fc Chimera Protein (4 µg/mL, R&D Systems). Chambered slides were mounted on a heated stage within a temperature-controlled chamber and maintained at 37°C and constant 5% CO2. The TIRFM set up acquisition was based on an Eclipse Ti2-E inverted microscope (Nikon) equipped with a 100×/1.45 NA Plan Apochromat LBDA objective (Nikon Instruments) and an iLAS 2 illumination control system (Roper Scientific SAS). Excitation was from a diode laser at 488nm (150 mW) (Vortran), 561nm (150 mW) (Coherent) and 642nm (110 mW) (Vortran) that passed a ZET405/488/561/647x filter (Chroma Technology). The emissions were separated and optically filtered by a ZT405/488/561/647rpc-UF1 dichroic (Chroma Technology) and ET525/50nm filter, ET610/75m filter, ET655lp filter (Chroma Technology) respectively. Images were recorded on a Prime 95B Scientific CMOS Camera or a Prime BSI Scientific CMOS Camera (Teledyne Photometrics). The final pixel size was 110 nm or 65 nm respectively. Image acquisition was controlled by MetaMorph Software (Version 7.10.5.476, Molecular Devices) and by Modular V2.0 GATACA software. The interval of time for acquisition was 3s – 5s.

For the double actin staining on conjugates, target cells were either un-pulsed or pulsed with 10 μM antigenic peptide (CMV p65) for 2 h at 37°C in RPMI 5% FCS/HEPES. Target cells were loaded 30 minutes with CellMask™ Green Actin Tracking Stain (5X) (Invitrogen™), washed three times and seeded at 2 × 10^4^ into µ-Slide 15 Well 3D glass bottom slides (Ibidi) coated with poly-D-lysine (Sigma). CTLs were loaded overnight with Wheat Germ Agglutinin Alexa Fluor 405 (1µg/mL) or Alexa Fluor 647 (0.5µg/mL) and loaded 30 minutes at 37°C with CellMask™ Orange Actin Tracking Stain (Invitrogen™), washed and seeded at 4 × 10^4^ per well. After 45 min of co-incubation at 37°C and constant 5% CO2 concentration, cells were fixed with 3% paraformaldehyde, washed with PBS/3% BSA/HEPES and directly acquired with a Yokogawa CSU-W1 Spinning Disk equipped of a Live Super-Resolution module, a ×100 Plan Apochromat LBDA objective (1.45 oil) (Nikon Instruments) and a Prime 95B Scientific CMOS Camera (Teledyne Photometrics).

### 3D-live spinning-disk microscopy

The 3D live imaging was performed on human CTL loaded overnight with Wheat Germ Agglutinin Alexa Fluor 488 (50 ng/mL, Molecular Probes, Invitrogen), and loaded 30 minutes with Vybrant™ DiD (1 µM) (Molecular Probes, Invitrogen). CTLs were seeded on µ-Slide 15 Well 3D glass bottom slides (Ibidi) coated with poly-D-lysine (Sigma), human monoclonal anti-CD3 antibody (1 µg/mL, TR66, Enzo Life Sciences) and recombinant human ICAM-1/CD54 Fc Chimera Protein (4 µg/mL, R&D Systems). Chambered slides were mounted on a heated stage within a temperature-controlled chamber maintained at 37°C and constant 5% CO2. During acquisition, the cells were in RPMI 1640 medium (1X) w/o pH Red supplemented with 10 mM L-Glutamine, 10mM Hepes and 5% fetal calf serum (FCS). The set up acquisition based on an Eclipse Ti2-E inverted microscope (Nikon) equipped with a 100×/1.45 NA Plan Apochromat LBDA objective (Nikon Instruments) and a Yokogawa CSU-W1 Spinning Disk. Excitation was from a diode laser at 488nm (150 mW) and 642nm (110 mW) (Vortran) that passed a ZET405/488/561/647x filter (Chroma Technology). The emissions were separated and optically filtered by a ZT405/488/561/647rpc-UF1 dichroic (Chroma Technology) and ZET405/488/561/647m (Chroma Technology). Images were recorded on a Prime 95B Scientific CMOS Camera (Teledyne Photometrics). The final pixel size was 110 nm. Image acquisition was controlled by MetaMorph Software (Version 7.10.5.476, Molecular Devices) and by Modular V2.0 GATACA software, the acquisition set up was on stream mode. The interval of time for acquisition was 2,48 - 3,48 seconds per z-stack.

### Structured illumination microscopy (SIM)

For high-resolution SIM (ELYRA PS.1; Carl Zeiss Microscopy) of the ultrathin sections, images were acquired by using the 63× Plan-Apochromat (NA 1.4) objective with excitation light of 405-, 488-, 561- and 642-nm wavelengths to visualize DAPI, WGA, GzmB and CellMask Deep Red, respectively. The DAPI image was recorded to identify both the nucleus of the CTL and the image plane. In a z-stack analysis 3-10 images were recorded with a step size of 100 nm to scan the cells of interest. ZEN 2012 software (Zeiss) was used for data acquisition and image processing to achieve higher resolution.

### Post-embedding correlative fluorescence electron microscopy (CLEM)

Transfected clonal CTLs in contact with JY cells were fixed with 3% PFA in D-PBS after different time points as described and kept at 4°C until high pressure freezing. For cryo-fixation cells were suspended in a 2% gelatin solution in D-PBS, supplemented with CellMask™ Deep Red (1:700, Invitrogen). Cells were dropped on sapphire discs coated with poly-L-orinithine (0.1 mg/ml) in flat specimen carriers (Leica). After incubation at RT (20°C ± 2°C) for 15 min to allow the cells to settle on the sapphire discs, the samples were vitrified in a high-pressure freezing system (Leica EM PACT2). Samples were further processed in a freeze-substitution system (AFS2; Leica) as described in ((*4*)). Briefly, all samples were transferred to the precooled (−130°C) freeze-substitution chamber of the AFS2. The temperature was increased from −130 to −90°C over 2 h. Freeze substitution was performed from −90 to −70°C over 20 h in anhydrous acetone and from −70 to −60°C over 24 h with 0.3% (w/v) uranyl acetate in anhydrous acetone. At −60 °C samples were infiltrated with increasing concentrations (30, 60, and 100% for 1 h each) of Lowicryl (3:1 K11M/HM20 mixture; Electron Microsopy Sciences). After 5 h of 100% Lowicryl infiltration, samples were UV polymerized at −60°C for 24 h and for additional 15 h at a linear temperature increase to 5°C. Samples were stored in the dark at 4°C until further processing. After removal of carriers and sapphire discs ultrathin sections (100 nm) were cut using an UC7 (Leica) and collected on carbon-coated 200 mesh copper grids (Plano). 1 d after sectioning the grids were stained with DAPI for 5 min (1/10,000), washed and sealed between a coverslip and a slide for high-resolution SIM imaging. After SIM imaging the analyzed grids were removed from the coverslips stained with uranyl acetate and lead citrate and recorded with the Tecnai 12 Biotwin electron microscope. Only CTLs with well-preserved membranes, cell organelles and nuclei were analyzed and used for correlation. For correlation, the DAPI (405 nm) and CellMask™ Deep Red (642 nm) signals, labeling the cell nucleus cell plasma membrane, respectively, were used to find the optimal overlay with the electron microscope images. The final alignment defines the position of the WGA 488 nm and GzmB-mCherry 561 nm fluorescent signals within the cell of interest. The Images were overlaid in Corel DRAW X6.

### GrzB-mCherry transfection of human CTLs

To monitor SMAPs secretion through cortical actin network 1 × 10^6^ human CTLs were pelleted in RPMI 1640 medium GlutaMAX (1X) with 10 mM Hepes, and washed once with Opti-MEM (Gibco). Cells were resuspended in 100 µL warm Opti-MEM with 2.5 µg capped and tailed RNA coding for GrzB-mCherry and transferred into an electroporation cuvette (0.2 cm, Bio-Rad) and electroporated with a Squarewave Gene Pulser Xcell Electroporation system (Bio-Rad), at 300 V for 2 ms. Cells were then seeded in 1 mL prewarmed RPMI/5%HS/human rIL-2/rIL15 medium at 37°C with 5% CO2 and used 16 hours after electroporation. Electroporated cells were loaded over night with WGA Alexa Fluor 405 (1µg/mL) or WGA Alexa Fluor 647 (0,1 µg/mL), washed and loaded 30 minutes with CellMask™ Green Actin Tracking Stain (4X) (Invitrogen™). Transfected and dye loaded CTLs were seeded on µ-Slide 15 Well 3D glass bottom slides (Ibidi) coated with poly-D-lysine (Sigma), human monoclonal anti-CD3 antibody (1 µg/mL) (TR66) (Enzo Life Sciences) and recombinant human ICAM-1/CD54 Fc Chimera Protein (4 µg/mL) (R&D Systems). CTLs were fixed with 3% paraformaldehyde, washed with PBS/3% BSA/HEPES and directly acquired with a Yokogawa CSU-W1 Spinning Disk equipped of a Live Super-Resolution module, a ×100 Plan Apochromat LBDA objective (1.45 oil) (Nikon Instruments) and a Prime 95B Scientific CMOS Camera (Teledyne Photometrics).

### 3D-LSR immunofluorescence on fixed CTL/Target cell conjugates

Target cells were either non-pulsed or pulsed with 10 μM antigenic peptide (CMV p65) for 2 h at 37°C and 5% CO_2_ in RPMI 5% FCS/HEPES, washed three times, and conjugated, using 1 minute centrifugation at 455g, with antigen-specific human CTLs previously loaded overnight with Wheat Germ Agglutinin (WGA) Alexa Fluor 647 (1 µg/mL) (Molecular Probes, Invitrogen). Conjugates were incubated for 2, 5, 15, 30 or 60 minutes at 37°C and 5% CO_2_ in RPMI 5% FCS/HEPES, gently disrupted, and subsequently seeded on Poly-L-Lysine-coated slides. Cells were fixed with 3% paraformaldehyde at 37°C for 10 minutes, washed with PBS/3%BSA/HEPES and permeabilized with 0.1% saponin (in PBS/3%BSA/HEPES). Conjugates were stained with the indicated primary antibodies (GzmB, clone GB11, #MA1-8073, Invitrogen; Lamp1, ab24170, abcam; CD107a, Clone H4A3, #555798, BD Biosciences, EEA1, clone F.43.1, #MA514794, Invitrogen; CD81, SN206-01, Invitrogen; Phalloidin-iFluor 405 Reagent, ab176752, abcam) for 1h at RT, washed 3x with PBS/3%BSA/HEPES/0.1% saponin and followed by isotype-matched secondary antibodies (Goat anti-rabbit IgG (H+L) Alexa Fluor ^TM^ Plus 405/488/555, Goat anti-mouse IgG1 Alexa Fluor ^TM^ Plus 488/555, Invitrogen) for 45 min at RT. Samples were washed and mounted in 90% glycerol-PBS containing 2.5% DABCO (Sigma) and examined using a Yokogawa CSU-W1 Spinning Disk equipped with a Live Super-Resolution module over a ×60 Plan Apochromat LBDA objective (1.40 oil) (Nikon Instruments) and a Prime 95B Scientific CMOS Camera. Z-Stack images of optical sections were acquired throughout the cell volume at 0.2μm intervals. Image acquisition was controlled by MetaMorph Software (Molecular Devices 7.10.5.476) and by Modular V2.0 GATACA software. Images were analyzed using ImageJ (<u>h,ps://imagej.net/ij/docs/faqs.html</u>) and quantified using Imaris Softare 10.2 (Oxford Instruments, Spot function and Shortest distance to surface func?on).

### Image analysis and quantification

Images were analysed using ImageJ (https://imagej.net/ij/docs/faqs.html).

Imaris Software 10.2 (Oxford Instruments) was used to measure SMAPs polarization towards the lytic synapse. Lytic synapse was manually defined using the surface model function on the phalloidin channel. Particles (WGA^+^, GzmB^+^ and WGA^+^GzmB^+^) were defined using the spots model function on granzyme B and WGA channels. Distances between each particle and the lytic synapse were measured.

To measure overlapping of WGA^+^ particles with EAA1, CD107a and CD81, each channel was defined using the surface model function in Imaris. The percentages of overlapping volume were calculated by measuring the sum of overlapped volume between WGA and EAA1 or CD107a or CD81 over the total volume of WGA.

### Reagents

| Target | Type | Supplier | Identifier | Dilution |
| --- | --- | --- | --- | --- |
| Granzyme B | Mouse IgG1 | Invitrogen | Clone GB11,<br>#MA1-80734 | 1/100 |
| Lamp1 | Rabbit mAb IgG | abcam | ab24170 | 1/100 |
| CD107a | Mouse IgG1 | BD Biosciences | Clone H4A3,<br>#555798 | 1/50 |
| EEA1 | Rabbit mAb IgG | Invitrogen | F.43.1,<br>#MA514794 | 1/20 |
| CD81 | Rabbit mAbIgG | Invitrogen | SN206-01 | 1µg/ml |
| Wheat germ agglutinin (WGA)<br>Alexa Fluor™ Plus 405 | Lectin | Invitrogen | #W56132 | 1µg/ml |
| Wheat germ agglutinin (WGA)<br>Alexa Fluor™ Plus 488 | Lectin | Invitrogen | #W11261 | 1µg/ml |
| Wheat germ agglutinin (WGA)<br>Alexa Fluor™ Plus 647 | Lectin | Invitrogen | #W32466 | 0.5-1µg/ml |
| Goat anti-rabbit IgG (H+L) Alexa<br>Fluor™ Plus 405 | Rabbit mAb IgG | Invitrogen | A48254 | 1/100 |
| Goat anti-rabbit IgG (H+L) Alexa<br>Fluor™ Plus 488 | Rabbit mAb IgG | Invitrogen | A32731 | 1/100 |
| Goat anti-rabbit IgG (H+L) Alexa Fluor™ Plus 555 | Rabbit mAb IgG | Invitrogen | A32732 | 1/100 |
| Goat anti-Mouse IgG1 Alexa Fluor™ 555 | Mouse IgG1 | Invitrogen | A21127 | 1/100 |
| Phalloidin-iFluor 405 Reagent | Recombinant Full Length Protein | abcam | Ab176752 | 1/50 |
| Human ICAM-1/CD54 Fc Chimera Protein, CF | human IgG1 | R&D System | #720-IC-2000 | 5µg/ml |
| Human monoclonal anti-CD3 | Mouse IgG1 | Enzo | TR66,<br>#ALX-804-822 | 10µg/ml |
| CellMask™ Green Actin Tracking Stain | Actin F | Invitrogen | #A57243 | 5x |
| CellMask™ Orange Actin Tracking Stain | Actin F | Invitrogen | #A57244 | 5x |
| Vybrant™ DiD Cell Labeling Solution | Fluorescent dye | Invitrogen | #V22887 | 1µM |
| Vybrant™ Dil Cell Labeling Solution | Fluorescent dye | Invitrogen | #V22885 | 1µM |
| SPY620-Actin | Actin F | Spirochrome | #SC402 | 4x |
| Poly-D-lysine-Hydrobromide | Adhesive | Sigma | #P6407 | 1:10 |
| Poly-L-lysine solution | Adhesive | Sigua | #P8920 | 1:10 |

---

**Supplementary Figure 1.**
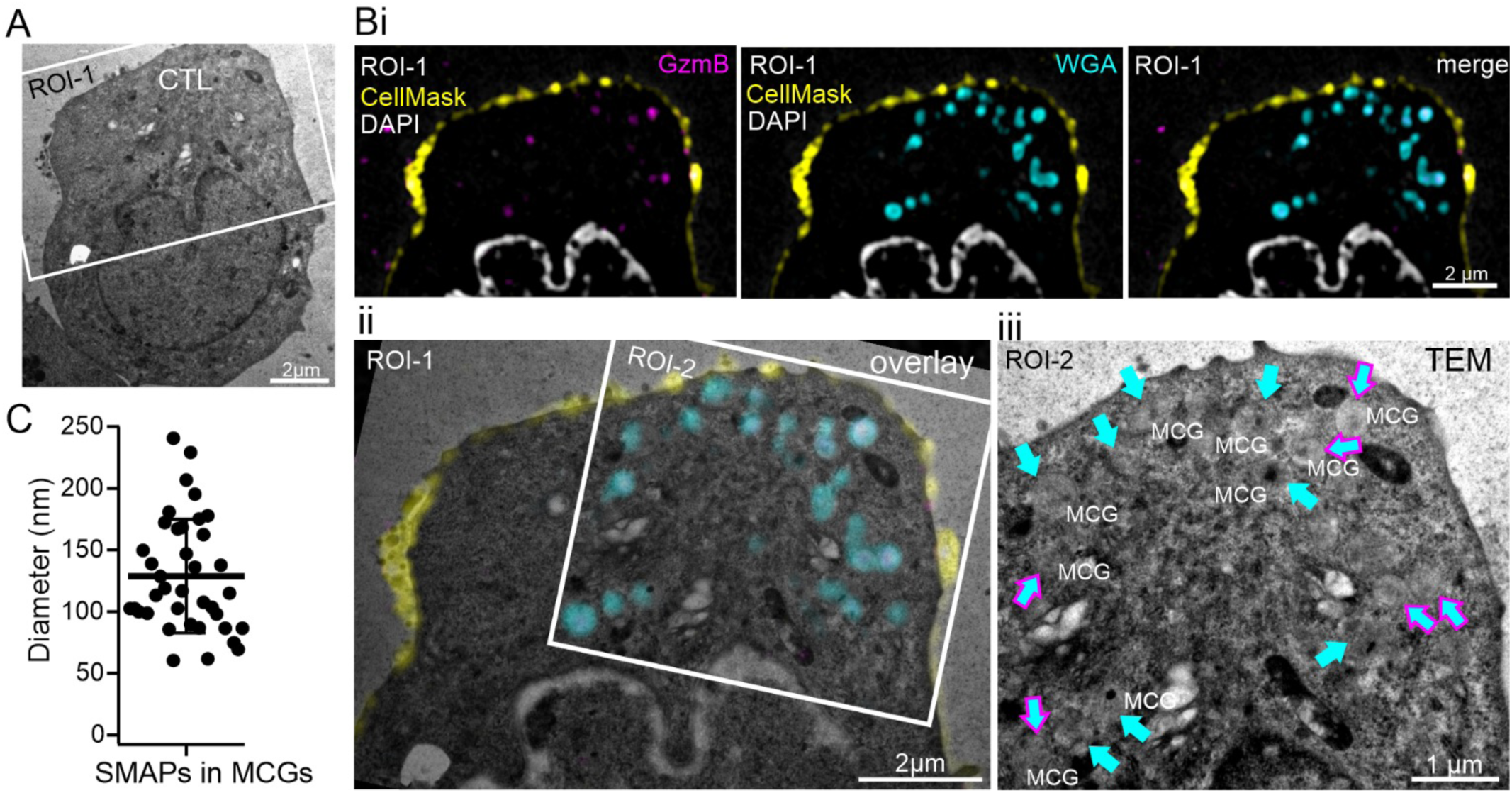
WGA^+^ and WGA^+^/GzmB^+^ particles are markers for MCGs. **(a)** Representative electron micrograph of a CTL without contact to a target cell. The white rectangle marks the area magnified in bi (ROI-1). **(bi)** Structured illumination microscopy (SIM) images of the magnified region (ROI-1 in A). The CTL was transfected with GzmB-mCherry and loaded with WGA-AF488. Shown are SIM images with GzmB (561 nm, magenta), WGA488 (cyan) and the merged signals. CellMask^TM^ DeepRed (yellow) and DAPI (white) fluorescence correspond to the plasma membrane and the nucleus, respectively. **(bii)** CLEM analysis of ROI-1 shown in (bi). The region marked with the white rectangle (ROI-2) is magnified in (biii). **(biii)** Magnified electron micrograph (ROI-2). Identified WGA^+^ and WGA^+^GzmB^+^ organelles are marked with arrows with the corresponding color code shown in the SIM images. MCG = multi core granule. **(c)** Diameter analysis of the SMAPs present in MCGs. Data are given as mean ± SD; N_MCGs_=23, n_SMAPs_=38.

**Supplementary Figure 2.**
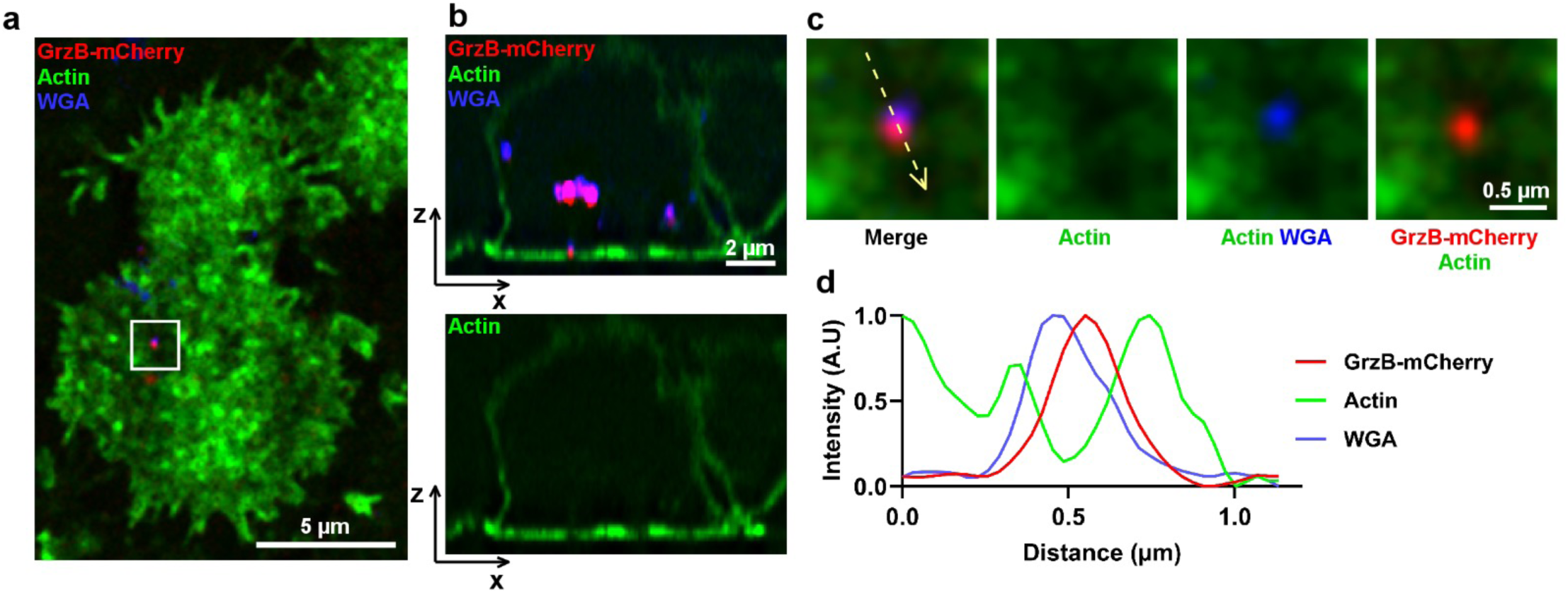
Secreted SMAPs percolate through the CTL actin cytoskeleton cortical network. **(a)** Shown is a structured illumination 3D spinning disk image of a representative CTL transfected with RNA coding for GrzB-mCherry (red), pre-loaded with WGA-AF647 (blue) and stained with CellMask^TM^ Green Actin (green). The white square marks the area magnified in (c). **(b)** x/z projection depicting a WGA^+^/GrzB^+^ SMAP particle crossing an area of actin clearance. **(c)** Magnification of the region delimited by the white square in (a) depicting a SMAP particle navigating through the actin cytoskeleton network. **(d)** Plot profile of the normalized intensity (arbitrary unit) of WGA, GrzB and actin along the yellow dashed arrows in (d). Data are from one representative experiment out 2.

**Supplementary Figure 3.**
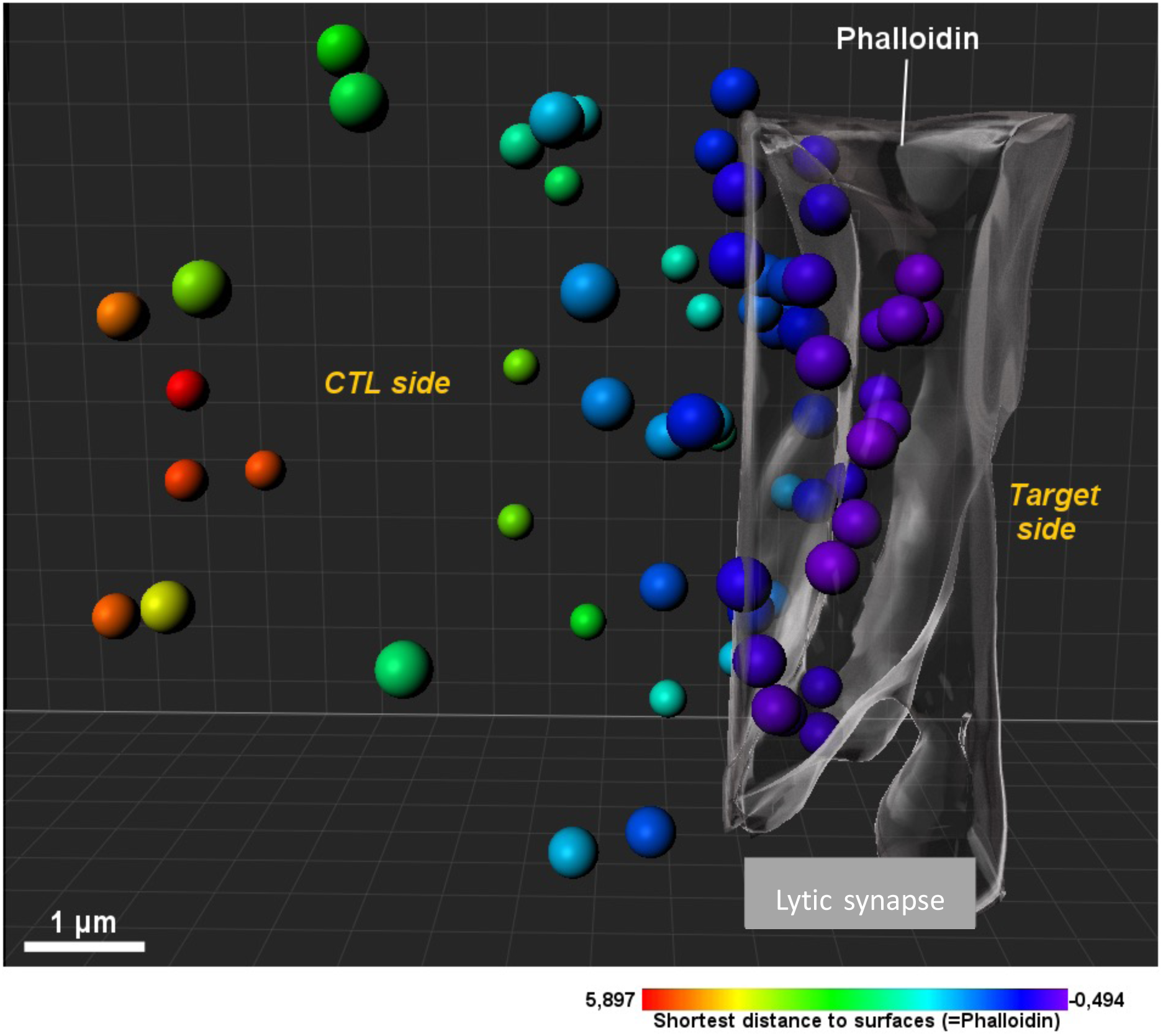
Schematic representation of distance to surface measurements using 3D-Imaris software. For all time points, 3D Imaris reconstructions of z-stacks acquired from CTL/target cell conjugates were used to measure the shortest distance to the cell surface (defined by cortical actin staining) for WGA^+^, GzmB^+^ and WGA^+^/GzmB^+^ particles. The pseudocolor scale indicates the shortest distance from the surface.

**Supplementary Figure 4.**
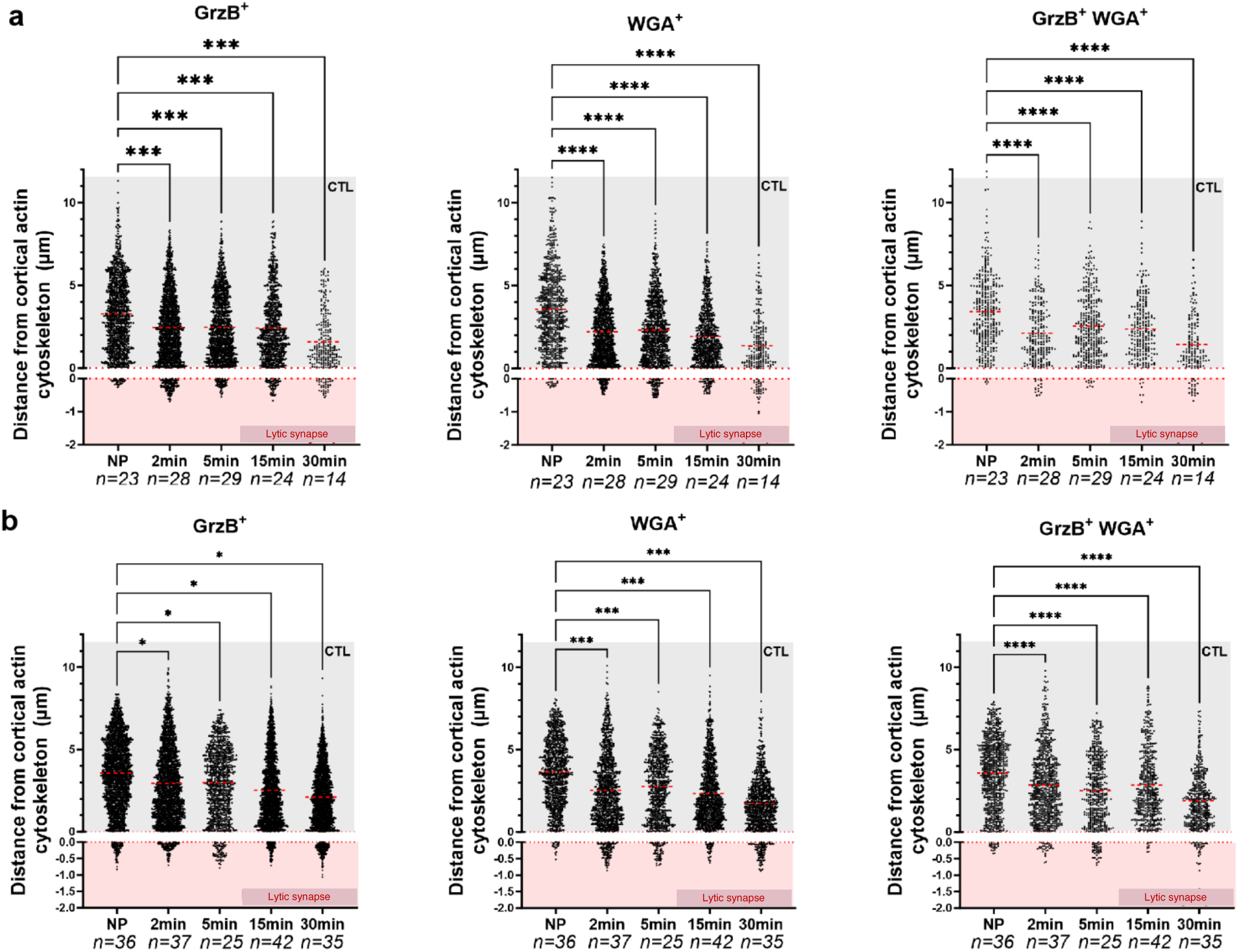
Time-dependent polarization of SMAPs towards lytic synapse. **a**) WGA^+^, GrzB^+^ and WGA^+^GrzB^+^ particle distances measured from synaptic cortical actin cytoskeleton over time in CTL /JY cell conjugates. Negative values indicate SMAPs detected within the cortical actin network of JY cells. (**b**) WGA^+^, GrzB^+^ and WGA^+^GrzB^+^ particle distances measured from synaptic cortical actin cytoskeleton over time in CTL /MDA-MB-231 cell conjugates. Negative values indicate SMAPs detected within the cortical actin network of MDA-MB-231 cells. Data are from 3 independent experiments for JY cells and 2 independent experiments for MDA-MB-231 cells. Statistical analyses were performed using GraphPad Prism 10.1.2. One-way ANOVA was utilized to assess statistical differences. \**p*<0.05, ** *p*<0.01, *** *p*<0.001, \*\*\*\**p*<0.0001, ns = not significant.

**Supplementary Figure 5.**
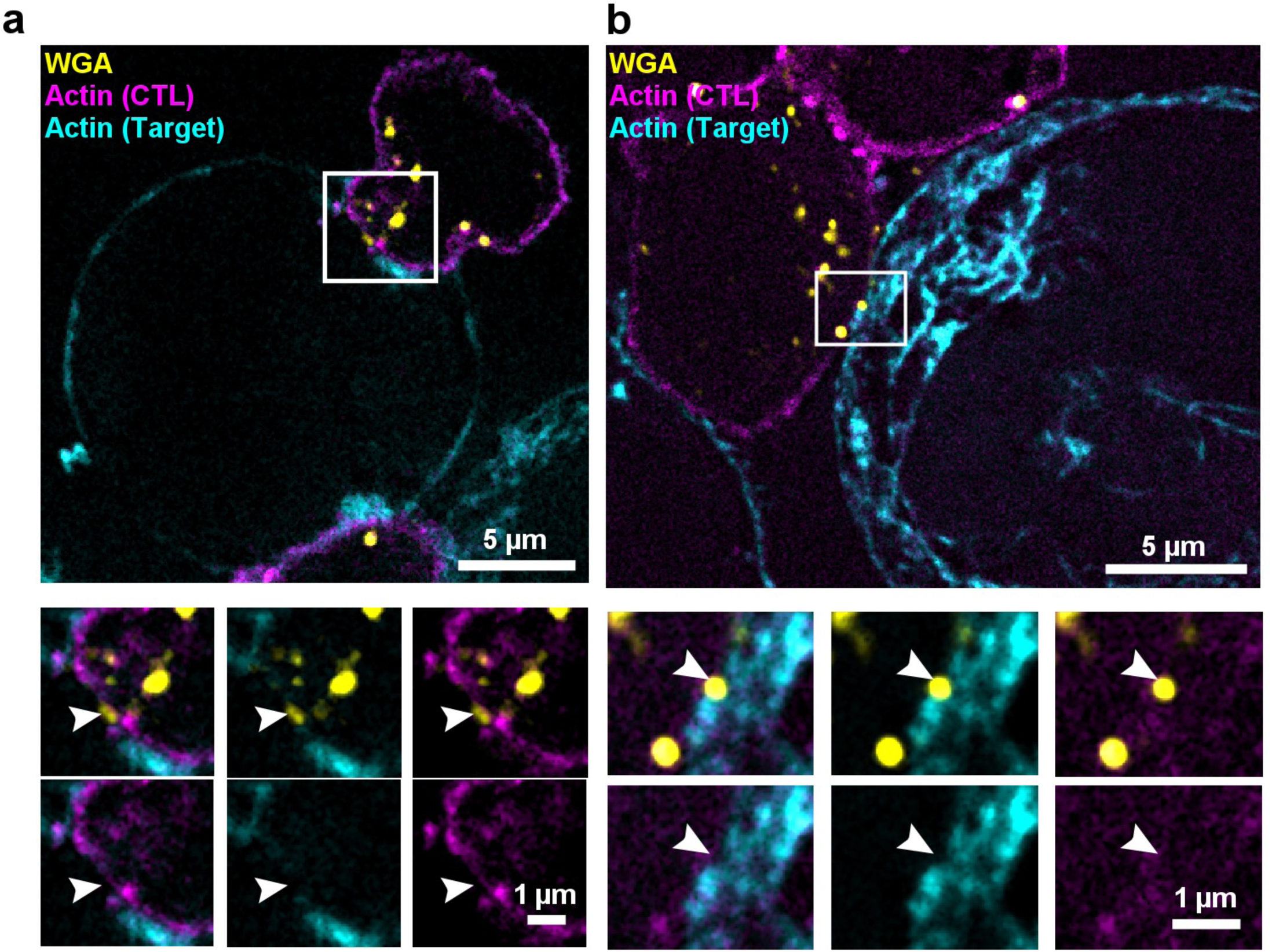
Intercellular SMAP transfer via *stenocytosis*. **(a** and **b)** Upper panels: Two representative conjugates formed between a CTL (previously loaded with CellMask^TM^ Orange Actin, magenta) and WGA-AF647 (yellow) and a target cell (JY) loaded with CellMask^TM^ Green Actin (cyan). The white rectangle marks the area magnified in lower panels. Lower panels: Magnification of the region delimited by the white square in (a and b). White arrows point the actin clearance areas in the CTL and in in the target cell. Data are from one representative experiment out 3.

**Supplementary Figure 6.**
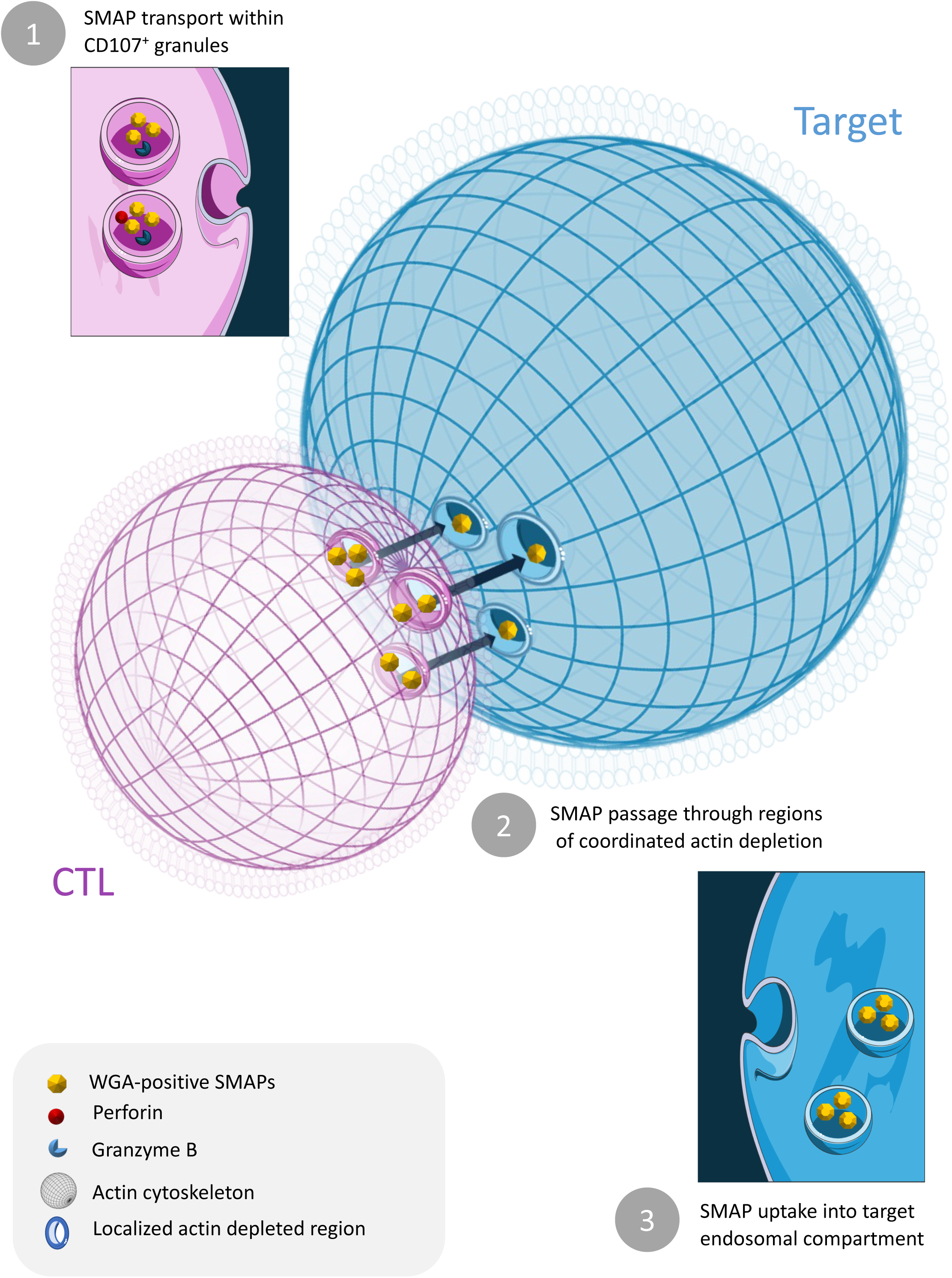
Schematic representation of *stenocytosis*. The drawing schematize schematizes stepwise transportation of SMAPs from CTL to target cells: 1) transport of SMAPs within CD107a⁺ granules in CTLs; 2) their passage through localized narrow regions of cortical actin depletion in both cells and 3) subsequent uptake into the target cell’s endosomal compartment.

## Movie legends

**Supplementary Movie 1**

Live TIRFM imaging (37°C/5%CO2) of a WGA-AF647 (pseudocolor) pre-loaded and Vibrant™ Dil (green) CTL seeded on an anti-CD3/ICAM-1 coated ibidi μ-slide chamber. Within minutes after stimulation, WGA^+^ immobile particles (pseudocolor) are detected in the TIRF plane in correspondence of areas exhibiting reduced plasma membrane staining (green).

**Supplementary Movie 2**

WGA^+^ particles only (pseudocolor) in the TIRF plane

**Supplementary Movie 3**

Areas exhibiting reduced Vibrant™ Dil plasma membrane staining (green)

**Supplementary Movie 4**

Live TIRFM imaging (37°C/5%CO2) of a WGA-AF647 (pseudocolor) pre-loaded and Vibrant™ Dil (green) CTL seeded on an anti-CD3/ICAM-1 coated ibidi μ-slide chamber. The cell is highly mobile with no detectable polarization of WGA-AF647^+^ particles (pseudocolor) in parallel with uniform Vibrant™ Dil (green) staining

**Supplementary Movie 5**

Ultra-rapid 3D Z-stacks acquisition of WGA-AF488 (magenta) preloaded CTLs stained with Vybrant™ DiD (cyan) and seeded on anti-CD3/ICAM-1 coated ibidi μ-slide chambers highlighting rapid docking and fusion events (cyan) of vesicles carrying WGA^+^ particles (magenta)

**Supplementary Movie 6**

Ultra-rapid Z-stacks acquisition of WGA-AF488 preloaded CTLs stained with Vybrant™ DiD (cyan) and seeded on CD3/ICAM-1 coated ibidi μ-slide chambers. Depicted are rapid membrane docking and fusion events (cyan).

**Supplementary Movie 7 and 8**

TIRFM imaging of WGA-AF488 (pseudocolor) preloaded CTLs stained with Vybrant™ Dil (green) and Actin SPY650 (white) and seeded on anti-CD3/ICAM-1 coated ibidi μ-slide chambers. Thinning out of the actin cytoskeleton network (white) overlap with hot-spots of SMAP (pseudocolor) release. Central cSMAC-like release patterns (**Supplementary Movie 7**) or patterns exhibiting multiple secretion-hot spots (**Supplementary Movie 8**).

**Supplementary Movie 9**

TIRFM imaging of a WGA-AF488 (pseudocolor) preloaded CTL stained with Vybrant™ Dil (green) and Actin SPY650 (white) and seeded on ICAM-1 only coated ibidi μ-slide chambers. No secretion patterns observed without anti-CD3 stimuli.

**Supplementary Movie 10**

Structured illumination 3D imaging confirms high transfection efficiency of GzmB-mCherry (red) together with WGA-AF647 staining (blue). Vybrant™ DiD (green) was used for membrane staining of transfected CTL cells.

**Supplementary Movie 11**

Spinning disk structured illumination z-stack acquisition of a conjugate between three WGA-AF647 preloaded CTL and an unpulsed target cell, fixed and stained with phalloidin (white) and antibodies against GzmB (green) and CD107a (red). Individual CTLs exhibit heterogeneity of cytotoxic content.

**Supplementary Movie 12**

3D reconstruction of the unpulsed conjugate from Movie 11 (see also Figure 4a). CD107a^+^ granules (red) contain single positive (WGA^+^ (green) and GzmB^+^ (blue)) and double positive (WGA^+^GzmB^+^) granules.

**Supplementary Movie 13**

Spinning disk structured illumination z-stack acquisition of a 2 minutes conjugate between a WGA-AF647 preloaded CTL and an peptide pulsed target cell, fixed and stained with phalloidin (white) and antibodies against GzmB (green) and CD107a (red). WGA^+^, GrzB^+^ and WGA^+^GrzB^+^ particles polarize towards the lytic synapse (see Figure 4b).

**Supplementary Movie 14**

Spinning disk structured illumination z-stack acquisition of a 30 minutes conjugate between a WGA-AF647 (green) preloaded CTL and an peptide pulsed target cell, fixed and stained with phalloidin (white) and antibodies against GzmB (blue) and CD107a (red). WGA^+^, GrzB^+^ and WGA^+^GrzB^+^ particles are detected inside the target cell (see Figure 4c).

**Supplementary Movie 15**

3D reconstruction of spinning disk structured illumination z-stack acquisition of a 15 minutes conjugate. WGA^+^ (green), GrzB^+^ (blue) and WGA^+^GrzB^+^ particles (SMAPs) are detected passing through nerrow gaps in the CTL/target cell cortical actin and taken up by target cells. Phalloidin (white, see Figure 7a).

